# Pancreas patch-seq links physiologic dysfunction in diabetes to single-cell transcriptomic phenotypes

**DOI:** 10.1101/555110

**Authors:** Joan Camunas-Soler, Xiaoqing Dai, Yan Hang, Austin Bautista, James Lyon, Kunimasa Suzuki, Seung K Kim, Stephen R Quake, Patrick E MacDonald

## Abstract

Pancreatic islet cells regulate glucose homeostasis through insulin and glucagon secretion; dysfunction of these cells leads to severe diseases like diabetes. Prior single-cell transcriptome studies have shown heterogeneous gene expression in major islet cell-types; however it remains challenging to reconcile this transcriptomic heterogeneity with observed islet cell functional variation. Here we achieved electrophysiological profiling and single-cell RNA sequencing in the same islet cell (pancreas patch-seq) thereby linking transcriptomic phenotypes to physiologic properties. We collected 1,369 cells from the pancreas of donors with or without diabetes and assessed function-gene expression networks. We identified a set of genes and pathways that drive functional heterogeneity in β-cells and used these to predict β-cell electrophysiology. We also report specific transcriptional programs that correlate with dysfunction in type 2 diabetes (T2D) and extend this approach to cryopreserved cells from donors with type 1 diabetes (T1D), generating a valuable resource for understanding islet cell heterogeneity in health and disease.

**Key findings:**

- Pancreas patch-seq provides a single-cell survey of function-transcriptome pairing in 1,369 islet cells from donors with and without diabetes
- Expression of a specific subset of genes predicts β-cell electrophysiology in transcriptome-function networks.
- Compromised β-cell function in T2D correlates with altered ETV1 expression and inflammatory pathways
- Functional heterogeneity in α-cells maps to ER stress and islet lineage markers
- Application of patch-seq to cells from rare cryopreserved islets from donors with T1D

## Introduction

Pancreatic islet cells control metabolism by releasing key hormones into the bloodstream via regulated exocytosis (Roscioni et al., 2016). For example, β-cells and α-cells respond to dynamic glucose levels by mounting electrical responses that culminate in regulated exocytosis of the two principal systemic glucoregulatory hormones, insulin and glucagon. These cells, especially insulin-producing β-cells, are long-known to show functional heterogeneity (Pipeleers, 1992). For instance, variations in β-cell insulin release, intracellular Ca^2+^ flux and electrophysiological activity have been described (Heimberg et al., 1993; Johnston et al., 2016; Salomon and Meda, 1986; Stefan et al., 1987; Zhang et al., 2003). Impaired β-cell function is a hallmark of the initial stages of type 2 diabetes (T2D) (Alejandro et al., 2015; Ashcroft and Rorsman, 2012; Kahn et al., 2006). Moreover, evidence suggests that β-cell function is regulated by paracrine action of other islet cell-types like γ- and δ-cells, although the mechanisms underlying this intra-islet regulation remain largely unresolved.

In parallel with known functional heterogeneity within islet cells, recent advances in single-cell technologies are uncovering molecular diversity across and within pancreatic cell-types at unprecedented molecular resolution (Tritschler et al., 2017). Single-cell RNA sequencing (scRNAseq) shows that the pancreas is transcriptionally diverse, revealing variable transcript enrichment across different islet cell-types and subpopulations (Baron et al., 2016; Enge et al., 2017; Muraro et al., 2016; Segerstolpe et al., 2016). This heterogeneity has been further refined through identification of surface markers and mass-spectrometry signatures (Bader et al., 2016; van der Meulen et al., 2017; Wang et al., 2016a). However, the ability to directly attribute islet cell molecular heterogeneity to physiologic properties, and functional deficits in disease, remains limited (Wang and Kaestner, 2018). For instance, the contribution of genes with altered expression in T2D (Segerstolpe et al., 2016) to functional consequences in islets remains unclear, and major gaps persist in our mechanistic understanding of T2D ‘risk’ candidate genes identified by GWAS (Mahajan et al., 2018; Prasad and Groop, 2016; Tritschler et al., 2017).

To address such limitations, we combined whole-cell patch-clamp measurements and scRNAseq (patch-seq) in dispersed islet cells, a method previously developed for neuronal studies (Cadwell et al., 2016; Földy et al., 2016; Fuzik et al., 2016). We collected 1,021 patch-seq cells from donors with no diabetes (ND) or from subjects with T2D, most of short duration, and dissected functional-gene expression relationships across islet cell-types using single-cell transcriptomes. Our data identifies genes and pathways driving functional heterogeneity in β-cells. We discovered a gene subset that predicts multiple β-cell functional phenotypes, and uncovered pathways driving increased exocytosis and ion-channel activity. Investigation of cells from T2D donors shows that key transcriptional regulators are enriched in T2D β-cells, in an apparent attempt to increase insulin secretion, but instead become drivers of β-cell dysfunction. Analysis of patch-seq alpha cells revealed heterogeneous expression of islet transcription factors and ER-stress genes that correlates to their electrophysiological profile. Finally, we extend this approach to 348 cells from rare islets of donors with T1D stored by cryopreservation, where we obtained profiles from α-, γ-, δ-, duct, and surviving β-cells. This work represents the first attempt to create a map of islet cell function-transcriptome pairing at the single-cell level and provides a valuable resource for exploring islet physiological and genetic dysfunction in diabetes.

## Results

### Pancreas patch-seq to simultaneously assess islet cell electrophysiology and transcriptome

To achieve patch-clamp followed by scRNAseq (patch-seq) in individual human islet cells, we initially isolated cells from 28 donors, including 18 with no diabetes (ND), 7 with type 2 diabetes (T2D), and 3 additional donors (**Supp Table S1**). We established pancreas patch-seq as a two-step process: (i) we performed electrophysiological measurements using standard whole-cell patch-clamp, and (ii) within 5 min from “break-in” we collected cellular content using a larger secondary pipette filled with lysis buffer (**Fig 1A**, Methods). This allowed intracellular access for whole-cell recording followed by recovery of full-length transcriptomes using SmartSeq2 that were sequenced to an average depth of 1-2 million reads (**Supp Fig 1, Methods**). A total of 1,021 cells (80%) passed quality control for both electrophysiology and sequencing, and were classified into major cell types based on the expression of key marker genes in a tSNE projection (**Fig 1B**, Methods). We obtained representatives of all major islet cell types (α-, β-, δ-, and γ-cells), and non-islet types such as acinar cells (**Fig 1C,D**). For each cell we measured parameters representing cell size, exocytosis, Na^+^ channel currents, and Ca^2+^ channel currents (**Fig 1E,F**). In this way, we obtained a broad survey of electrophysiological activity of all major islet cell-types in both ND and T2D settings (**Supp Fig 2**). In addition to expected cell-type differences in size measured by whole-cell capacitance (**Fig 1C,F**), these data demonstrate substantial variation in exocytosis and channel activity in different pancreatic cell types (**Supp Fig 2**).

**Figure 1.**
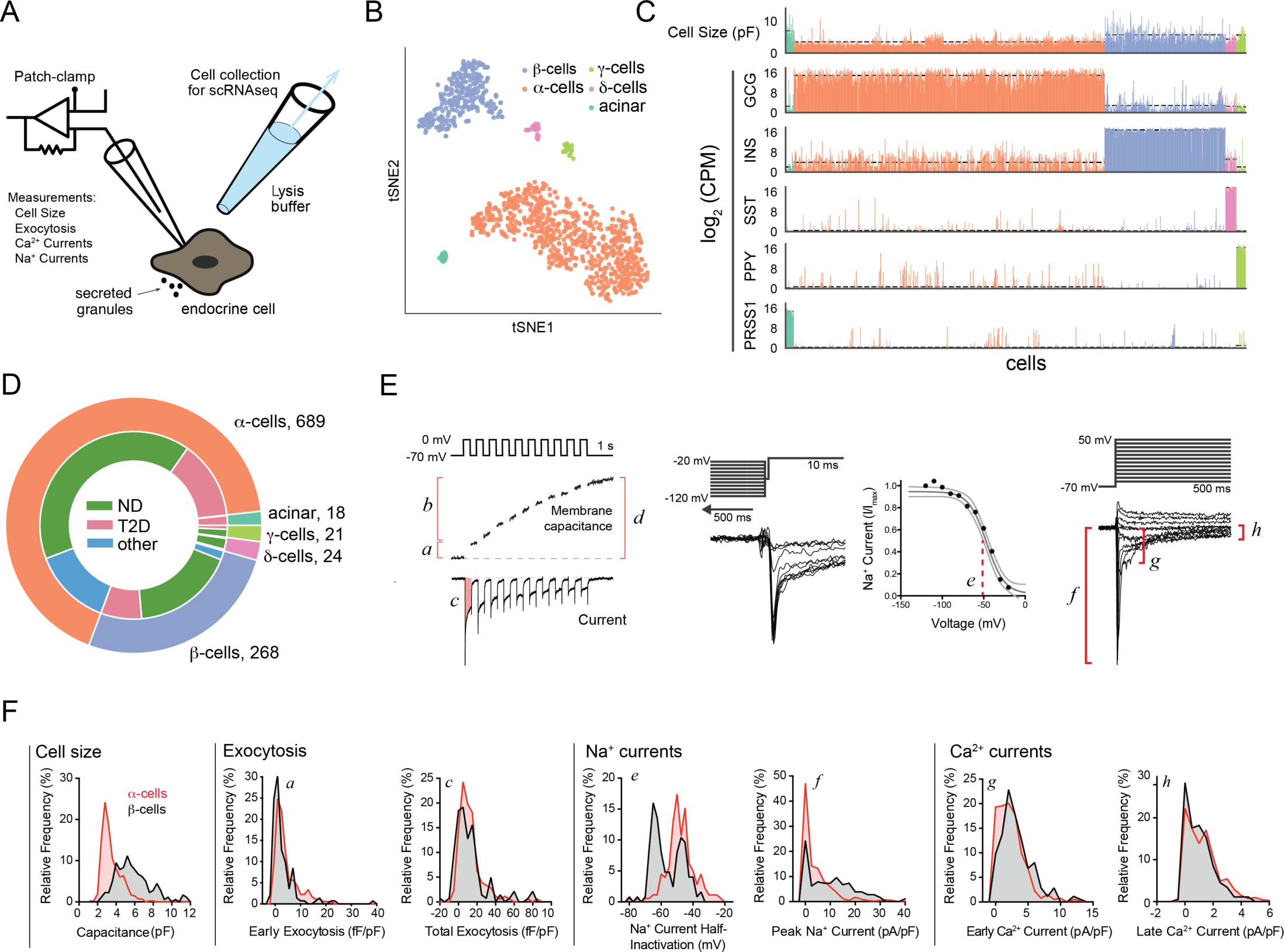
Pancreas patch-seq pipeline. (A) Schematic of pancreas patch-seq protocol. (B) tSNE projection of measured patch-seq cells clustered by gene expression of over-dispersed genes. Cells are colored by cell type according to cluster identification by marker genes. (C) Cell size measured as membrane capacitance for each cell compared to expression of key marker genes. Color indicates cell type according to color code in panel B. Dashed line shows average capacitance/gene expression for each cell type. (D) Distribution of total number of patch-seq cells based on cell-type and diabetes status. (E) In each cell we measured: *a*-early exocytosis; *b*-late exocytosis; *c*-Ca^2+^ integral, *d*-total exocytosis, *e*-Na^+^ current half-inactivation; *f*-peak Na^+^ current; *g*-early Ca^2+^ current; *h*-late Ca^2+^ current (not shown, we also measured cell size, reversal potential, and Na^+^ and Ca^2+^ conductance). (F) Distribution of selected parameters demonstrating heterogeneity in functional responses of α-(red) and β-(black) cells. Inset letters (*a-h*) correspond to measured parameters in panel E. Distribution of exocytotic responses in *a* and *c* are shown for 1 mM glucose (α-cells) or 5-10 mM glucose (β-cells).

To rigorously assess the robustness of our pancreas patch-seq pipeline, we also collected an additional 3,518 cells by FACS for scRNAseq *without* patch clamping from 14 of these donors (8 ND, 6 T2D; **Supp Fig 1** and Methods). The transcriptomes of cells after patch-seq or FACS-purification from the same donors led to comparable quality metrics (**Supp Fig 3**). While we observed a difference in the number of genes with detectable transcript, the values from cells after patch-seq or FACS were within the range of previous datasets (**Supp Fig 3**). Analysis of genes required to maintain fundamental cellular functions (hereafter, ‘housekeeping’), islet identity, and immediate early genes (IEG) showed that most differences are driven by (i) varying sensitivity to genes expressed at low levels, and (ii) varying expression of stress-response genes likely reflecting unavoidable steps like islet shipping and dispersion (**Supp Fig 3**). Moreover, we found that with longer islet culture, the initial increase in IEG transcripts decreased, while transcripts encoding islet-specific genes increased (**Supp Fig 4**). Overall, gene expression in patch-clamped cells was comparable to prior datasets (Segerstolpe et al., 2016), showing a high number of islet-specific transcripts and low IEG activation (**Supp Fig 3**). Finally, patch-seq itself did not appear to impact gene expression since tSNE data plots from cells collected without whole-cell patch-clamp, overlapped with the patch-clamped cells (**Supp Fig 4**).

### Pancreas patch-seq links transcriptional and functional heterogeneity of β cells

We applied the patch-seq approach to find genes associated with electrical and exocytotic function in β-cells, where the electrophysiological properties cluster by ‘functional group’ (exocytosis, Ca^2+^ and Na^+^ currents) and are uncorrelated with other parameters (**Fig 2A; Supp Fig 5**). We first correlated the transcriptome of β-cells to total exocytosis, which is representative of the total secretory capacity of a β-cell under stimulatory glucose levels, and a key indicator of β-cell function and dysfunction in T2D (Ferdaoussi et al., 2015; Gembal et al., 1992). In this way, we found genes positively or negatively associated with β-cell secretory capacity (**Fig 2B**). Among top correlates, we found several genes linked to pathways thought to regulate insulin secretion, including β-cell transcription factors (*MAFA, ETV1*), molecules associated with insulin granules (*SLC30A8, VAMP2, SCG2, INS*), metabolic enzymes (*PDK4, PDHA1, GYG1*), and ion channels (*ABCC9, KCNH2, KCNMB2, NALCN*) including the L-type Ca^2+^ channel encoded by *CACNA1C* (Rorsman and Braun, 2013; Lu et al., 2002; Ait-Lounis et al., 2010; Zhang et al., 2005; Chimienti et al., 2004). Gene set enrichment analysis (GSEA) using correlation scores confirmed many of these pathways and revealed additional enriched categories, such as neuronal regulators, transcription factors, and regulators of cell-polarity or stress (**Fig 2C**; **Supp Fig 5**). Islet transcription factors with weaker but significant association to exocytosis included *PAX6*, *FOXO1* and *NKX6-1* (**Supp Fig 5**).

**Figure 2.**
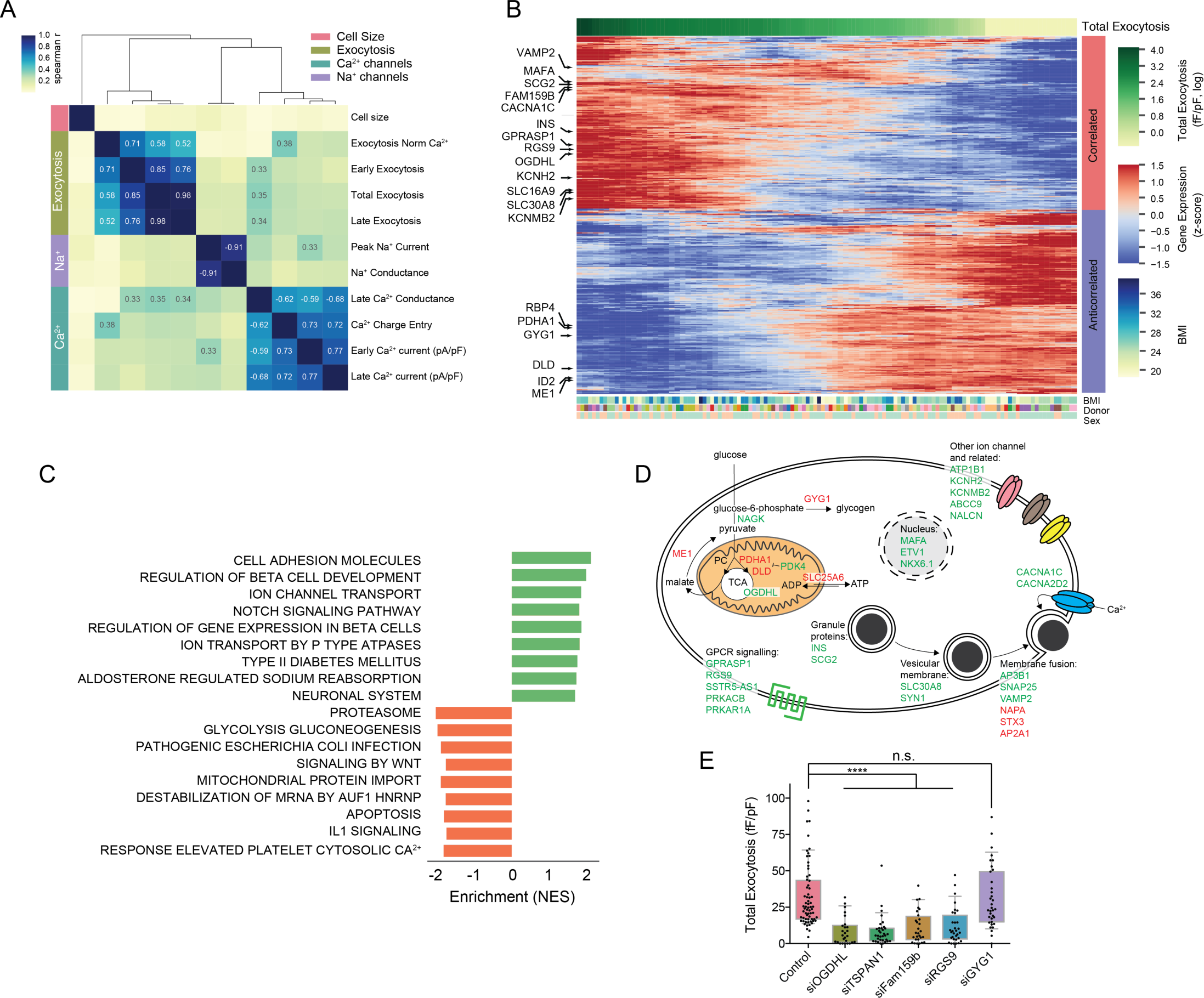
Correlation of β-cell exocytosis to single-cell gene expression and pathway analysis. (A) Spearman correlation of measured electrophysiological parameters shows clustering of each functional group (Exocytosis, Ca^2+^ currents, Na^+^ currents, Cell size) and low cross-correlation across clusters. All parameters are normalized to cell size. (B) Heatmap of top correlated and anticorrelated genes to measured total exocytosis in ND β-cells at 5-10 mM glucose. Cells are sorted by exocytotic response from highest (left, dark green) to lowest (right, yellow). Gene expression is shown as a z-score after smoothing (n=20 cells). Metadata associated to each cell (BMI, Donor, Sex) is shown at bottom. (C) Top enriched pathways in GSEA using genes correlated (red) and anticorrelated (blue) to total exocytosis (pathways in KEGG and Reactome databases, FDR<0.1). (D) Summarized map of cellular location and key pathways with genes correlated (green) and anticorrelated (red) to exocytosis. (E) Exocytosis in β-cells measured at 10 mM glucose following target gene knockdown compared to control cells from same donors (348 cells, N≥3 donors per knockdown experiment). ** FDR<0.01, **** FDR<0.0001 (Mann-Whitney-U test and BH correction).

The majority of genes whose expression correlated significantly with β-cell exocytosis are islet-specific and also included candidate T2D risk genes like *YWHAG* (Fernandez-Tajes et al., 2018). Top gene correlates to exocytosis also included regulators of oxidation and detoxification that could mitigate T2D-associated stress (Otter and Lammert, 2016), like glutathione peroxidase and transferases (*GPX3, GSTK1* and *GSTA4*: **Supp. Table S2**). We integrated these results into a map that combined correlations of transcript levels with four exocytosis measures (early, total, late, and normalized to Ca^2+^; **Fig 2D**), permitting visualization and analysis of gene expression significantly correlated with glucose-dependent insulin secretion in β-cells.

This analysis nominated multiple genes - without known roles in β-cells - as candidate regulators of physiological function, including *OGDHL, FAM159B, TSPAN1, RGS9* and *GYG1*. Prior studies have linked *OGDHL* to metabolism, while *FAM159B* and *TSPAN1* encode membrane proteins, and *RGS9* specifies a regulator of G-protein signalling (Bunik et al., 2008; Danielsson et al., 2014; Uhlen et al., 2015). To test our predictions, we performed siRNA knock-down followed by patch-clamp for this set of genes in islets from an additional 8 ND donors (**Supp Table S3; Fig 2E**). Knockdown of each gene, confirmed by qPCR (**Supp Table S4**), was followed by electrophysiological characterization at 10 mM glucose and insulin immunostaining to confirm cell-type. In 4/4 cases of genes positively correlated to exocytosis (*OGDHL, FAM159B, TSPAN1* and *RGS9*), we observed significant reduction of the exocytotic responses after knock-down (**Fig 2E**). By contrast, knock-down of *GYG1*, whose transcript levels are anticorrelated with exocytosis, did not lead to significant changes of exocytosis, possibly reflecting that β-cell activity is already maximally-stimulated at these high-glucose conditions. Together, these data show that patch-seq analysis robustly identified unsuspected genetic regulators of human β-cell physiology.

### Patch-seq gene expression networks that predict β-cell electrophysiological phenotypes

To obtain further insight about genes linked by patch-seq to β-cell excitability, we extended our analysis to additional electrophysiological parameters (Ca^2+^ currents, Na^+^ currents), and selected transcripts showing consistent positive or negative correlations to multiple parameters as likely regulators of β-cell function (**Fig 3A**). This analysis identified several key genes, including genes previously associated with β-cell transcriptional heterogeneity (Avrahami et al., 2017; Dorrell et al., 2016; Segerstolpe et al., 2016; Tritschler et al., 2017). For instance, we observed transcripts encoding retinol binding protein (*RBP4*) correlated negatively with cell size, Na^+^ currents and total exocytosis, while transcripts encoding the β-cell surface protein *FAM159B* correlated positively to exocytosis, Ca^2+^ entry and Na^+^ currents. A tSNE projection using these highly correlated genes shows a gradient of functional measures across β-cells that overlaps with patterns of gene expression (**Fig 3B**). To understand further how genetic pathways drive each major group of physiological function (exocytosis, Ca^2+^ currents, and Na^+^ currents), we repeated GSEA on their averaged correlation scores (**Supp. Fig 6**). Results for overall exocytosis are consistent with those reported above (**Fig. 2C**); by contrast, GSEA further identified that Na^+^ and Ca^2+^ currents are linked to pathways related to increased excitability and circadian rhythms respectively, consistent with prior studies (Perelis et al., 2015). Several of the observed pathways identified by GSEA on averaged correlation scores also overlap with those recently implicated in islet dysfunction in T2D using genomic data (Fernandez-Tajes et al., 2018)

**Figure 3.**
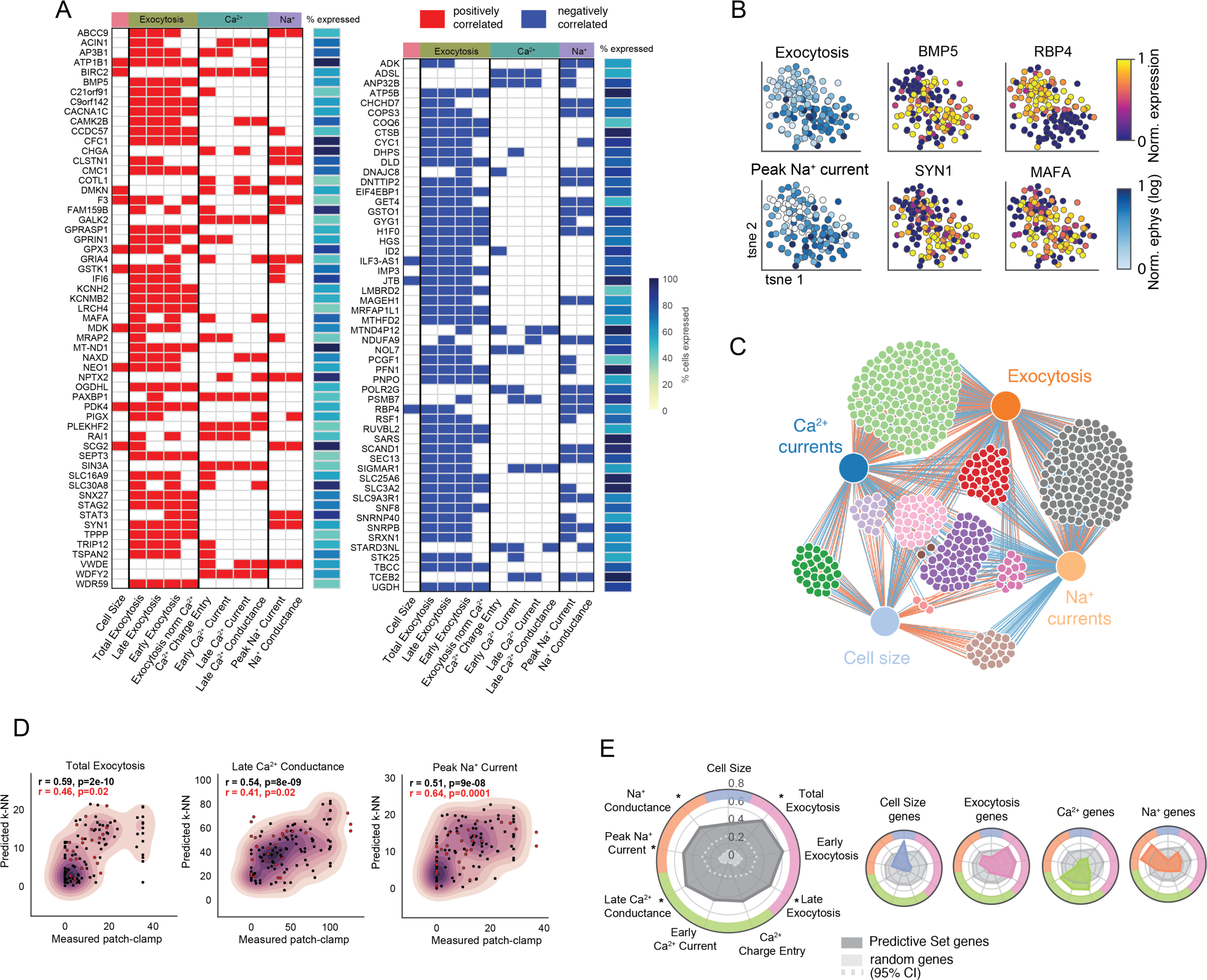
Gene networks driving β-cell functional heterogeneity. (A) Genes showing significant correlations to several electrophysiological parameters (z-score>2 for N>5 parameters, see Methods). Significant positive/negative correlations are indicated in red/blue. (B) tSNE projection of ND β-cells using genes discovered in panel A. Color indicates normalized expression of each gene (MAFA, SLC30A8, RBP4, ID2) or functional parameter (Total exocytosis, Peak Na^+^ current). (C) Network of genes connecting different functional groups (‘PS genes’). Genes (small dots) connecting different functional groups (large dots) are selected if they show significant correlations (z-score>2) to at least one functional parameter in each group. Edge color indicates positive/negative correlations (red/blue), and gene color identifies clusters connected to the same functional groups. (E) Prediction of β-cell electrophysiology from gene expression. Representative plots of measured patch-clamp values versus predicted values from nearest neighbours in gene expression using ‘PS genes’ (k-NN model, n=5) for a training (black) and validation (red) dataset. Spearman correlation and p-value for each parameter are indicated as inset. (F) Spider plot showing performance of the model using the ‘PS genes’ (dark gray), measured as Spearman correlation between experimental and predicted values. Comparison to mean values using 484 random genes of equivalent expression in β-cells (10,000 permutations, 95% CI interval shown in light gray). Asterisk indicates parameters for which we also obtained significance (p<0.05) in a validation set of data withheld from the correlation analysis. Smaller spider plots show performance of the ‘PS gene set’ (gray) versus top correlated genes for each functional group: exocytosis (pink), Ca^2+^ (green), Na^+^ (orange), cell size (blue).

We then asked whether genes that correlate to multiple groups of physiological function (“functional groups”) could be used to develop predictive algorithms that link transcriptional signatures to β-cell function. To do so, we generated a network of genes with significant correlation to more than one functional group (e.g. Ca^2+^ and exocytosis), and selected those with highest median expression (**Fig 3C, Supp Table S5; Supp Material S1**). This list of highly-connected genes, that we termed our Predictive Set (PS) contains genes: (i) previously linked to β-cell excitability, (ii) with heterogenous expression in β-cell subpopulations and in T2D (FFAR4, FXYD2, ID4), and (iii) with unknown function in β-cells (**Supp Table S5**). We then attempted to predict the measured electrophysiology of each cell from the average values of its nearest neighbours (NN) in PS gene expression (N=484 genes, k-NN with 5 neighbors) (see Methods). We obtained significant correlations between the experimentally measured patch-clamp parameters and the predictions from the k-NN model (**Fig 3D; Supp Fig 7, black**). We compared the performance of PS genes to a list of randomly selected genes of equivalent expression in β-cells (N=10,000 permutations), and found that the PS genes performed significantly better for all parameters (**Fig 3E**). We also tested our model by comparing its predictive power to that obtained using the top genes correlated to each functional group; finding that PS genes showed a consistent performance across all measured parameters, whereas group-specific gene sets only performed well for their subset of parameters (**Fig 3E**). Finally, we validated the predictive power of PS genes with a ‘test’ set of data including gene expression and functional parameters of cells withheld from the analysis that generated the PS gene set. This analysis demonstrated appropriate recovery of significant correlations for 5 of the measured functional parameters **(Fig 3D; Supp Fig 7, red, and Fig 3E**). Thus, patch-seq and network analysis combined with machine learning generated unprecedented algorithms that reliably link gene expression to global β-cell function.

### Markers of β-cell heterogeneity correlate to β-cell excitability

Within the identified ‘PS gene set’ we found that transcripts encoding RBP4 are significantly correlated with β-cell functional heterogeneity. Prior studies have noted heterogeneous expression of *RBP4* in β-cell subsets, but did not establish direct links to functional heterogeneity (Baron et al., 2016; Dorrell et al., 2016; Rui et al., 2017; Segerstolpe et al., 2016). Other studies suggest that RBP4 is an adipokine with roles in homeostatic regulation of metabolism (Broch et al., 2007; Tritschler et al., 2017). We observe that *RBP4* transcript levels have significant correlation with multiple β-cell functional phenotypes; for example, it is the ‘PS gene’ with strongest anti-correlation to β-cell Na^+^ channel activity. RBP4^+^ β-cells have decreased exocytosis and Na^+^ currents despite having normal Ca^2+^ current activity (**Supp Fig 8**). Consistent with this, we observed that RBP4^+^ β-cells also had significantly reduced expression of key regulators of β-cell stimulus-secretion coupling like *KCNJ8*, *ABCC9* and *SCN3A* (**Supp Fig 8**); this latter gene encodes the principal physiologically-relevant Na^+^ channel in rodent β-cells (Zhang et al., 2014). Together, these results provide evidence that *RBP4* expression is a marker of β-cells with reduced function, and show that some previously-identified markers of β-cell transcriptomic heterogeneity identify subpopulations with heterogeneous function (Dorrell et al., 2016; Johnston et al., 2016).

### Impaired function and gene expression in β-cells from donors with T2D

Pancreas patch-seq in islets from the 7 donors with T2D (**Supp Table 1**) showed reduced β-cell insulin content and secretion (**Fig 4A**), and exocytosis (**Fig 4B, Supp Fig 9**). Except for one donor (R241, who did not contribute β-cells for analysis: see Methods), all β-cells were obtained from donors with recent T2D onset (<9 years), and who were not receiving insulin treatment. To identify genetic drivers of T2D β-cell dysfunction, we assessed transcripts correlated to exocytosis in ND controls (**Fig 2B, Supp Table 2**). We observed that genes that positively correlate with exocytosis in ND β-cells are *up-regulated* in T2D, while genes that negatively correlate with exocytosis in ND β-cells had *reduced* expression in T2D (**Fig 4C**). These data suggest that β-cells attempt to alter transcript levels in T2D, likely in response to increased insulin demand, but ultimately fail to compensate, since their exocytotic response is impaired compared to ND controls (**Fig 4B**).

**Figure 4.**
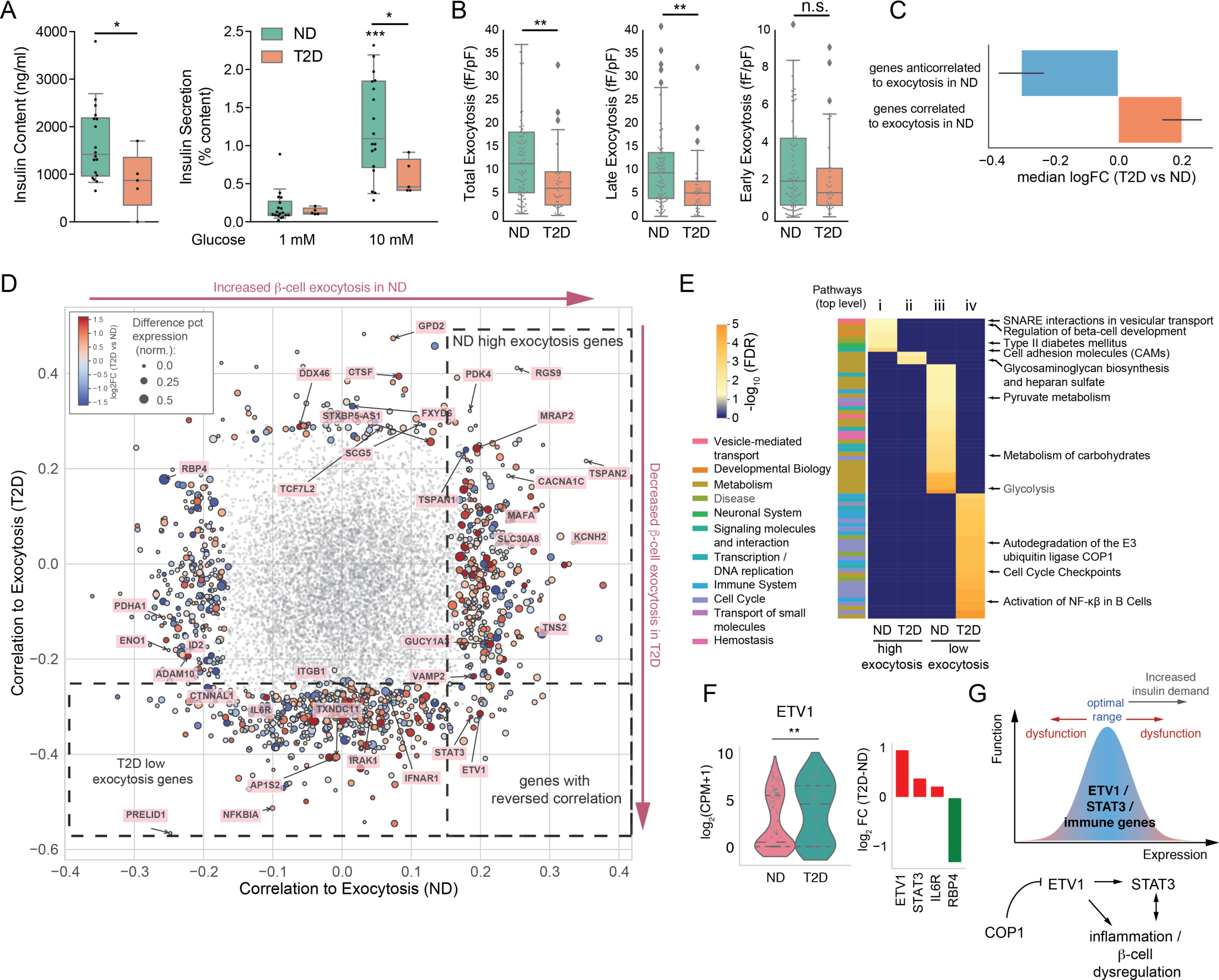
Functional and transcriptomic changes in β-cells early in T2D. (A) Insulin content and secretion (as % content) for donors used in this study. *p<0.05, ***p<0.001 (Student’s t-test; Two-way ANOVA and Tukey post-test) (B) Distribution of measured exocytosis parameters for ND and T2D β-cells. ** FDR<0.01 (Mann-Whitney-U test and BH correction). (C) Variation in T2D of genes found to be correlated (red) or anticorrelated (blue) to exocytosis in nondiabetic cells. Data is shown as median log fold-change in expression between T2D and ND. Error is SEM. (D) Gene correlation map of exocytosis. Scatter plot shows correlation to exocytosis in ND (x-axis) and T2D (y-axis) for each gene. Genes with significant correlations (z-score>2) are colored according to their fold-enrichment in T2D cells (red enriched in T2D, blue enriched in ND), and size is proportional to change in % expression (larger dots have different % of cells expressing the gene in T2D and ND). Regions of interest are highlighted with dotted boxes. Genes with non-significant correlations in gray. (E) Enriched pathways for genes correlated to exocytosis in ND (i), T2D (ii) and genes anticorrelated to exocytosis in ND (iii), T2D (iv). Enrichment is shown as log_10_(FDR) and blue indicates that the pathway is not enriched for a given set. Left bar indicates top category of each enriched pathway. (F) Distribution of ETV1 expression (left) and log enrichment of a subset of genes between ND and T2D in the patch-seq dataset. (G) Model showing the hypothesized role of ETV1, STAT3 and immune pathways in β-cell dysfunction in early T2D. Based on (Suriben et al., 2015).

To find genes that could underlie this effect, we performed correlation analysis in T2D cells, and integrated the ND and T2D results in a correlation map of exocytosis with the overall changes in gene expression (**Fig 4D, Supp Table 2**). Next, we performed a pathway analysis for each of the 4 gene subsets (ND/T2D and correlated/anticorrelated to exocytosis) (**Fig 4E**, **Supp Fig 10)**. Whereas low exocytosis in ND β-cells is linked mostly to metabolic pathways, dysfunction in T2D is related to immune response, cell cycle pathways and altered transcription factor expression. In particular, we observe induction of pathways like NFκB signalling, and auto-degradation of the ubiquitin ligase COP1. Genes in T2D that show increased expression associated with reduced β-cell exocytosis included *NFKBIA, IL6R* and *IRAK1*, known to encode regulators of immune and inflammatory pathway, and the transcription factors *ETV1* and *STAT3* (**Fig 4E-F, Supp Fig 10**). The ETV transcription factors impair insulin secretion in hyperglycaemic mouse models, and are negatively regulated through COP1–dependent targeted degradation (**Fig. 4G**) (Suriben et al., 2015). ETV1 shows negative correlation to several measured exocytosis parameters in T2D (**Supp Fig 10**). We tested genes previously reported to show ETV-dependent expression in mouse islets (Suriben et al., 2015), and found several human orthologs in our β-cell single-cell correlations, including *STAT3, MAFA, SLC30A8*. Analysis of these ETV-dependent genes showed an enrichment of (i) pathways that we found to be important for β-cell exocytosis in our correlation analysis, and (ii) genes related to immune cytokine signaling, like *STAT3* which encodes a critical cytokine signaling factor, that also has postulated roles in insulin secretion, β-cell regeneration and neonatal diabetes (O’Shea and Plenge, 2012; Saarimäki-Vire et al., 2017). Finally, we analyzed full-length transcriptomes to investigate ETV1 splicing in β-cells and detected no signs of differential splicing between ND and T2D (**Supp Fig 10**). These results indicate that expression of transcriptional regulators and their downstream targets requires balance for optimal insulin secretion, and that an imbalance may lead to dysregulated secretion during early T2D (**Fig 4G)**.

### Transcriptional and physiological heterogeneity observed in α-cells

Cell size and Na^+^ channel properties are often used to identify α-and β-cells in rodents (Briant et al., 2017; Zhang et al., 2014); however this is not generally applicable to humans, making it difficult to distinguish islet cell types at the time of patch-clamping. For example, α-cells (definitively identified *post hoc* from scRNAseq) were frequently mis-classified as possible β-cells during initial patch-clamp measures of capacitance that indicated cell size (>4 pF; **Fig 1F**). To improve cell-type identification prior to scRNAseq, we implemented a machine learning algorithm based in ‘random forests’ (Breiman, 2001) to classify cell types from their electrophysiological fingerprint. We trained and validated the model in cells from ND donors using 10-fold cross-validation, obtaining an AUC of 0.95 in the validation dataset (**Fig 5A,B**). Features from every functional group contributed to the classifier (**Fig 5C**), and the model also performed well in cells from T2D donors (AUC=0.93) (**Fig 5A**). Compared with a capacitance-based cell size cut-off at the time of patch-clamping, our model significantly improved the *a priori* identification of α-cells, and reduced mis-classification as β-cells (**Fig 5D**). This advance improved investigation of α-cell transcriptomes and revealed significant transcriptional heterogeneity in islets from multiple human donors, with a subset of cells showing an enrichment in genes related to α-cell maturation (LOXL4, MAFB, ARX), evidence of reduced ER stress (DDIT3, XBP1, PPP1R15A), transcription factors governing endocrine fate (FEV, ISL1), receptors involved in glucose homeostasis (FFAR1, GPAR119), and ion channels (KCNK16) **(Fig 5D; Supp Fig 11**). This transcriptional heterogeneity correlated to electrophysiological features including Na^+^ currents and cell size (**Fig 5D**). Thus, in addition to improving prospective identification of patch-clamped human α-cells, our studies provide evidence for molecular heterogeneity underlying functional heterogeneity in α-cells.

**Figure 5.**
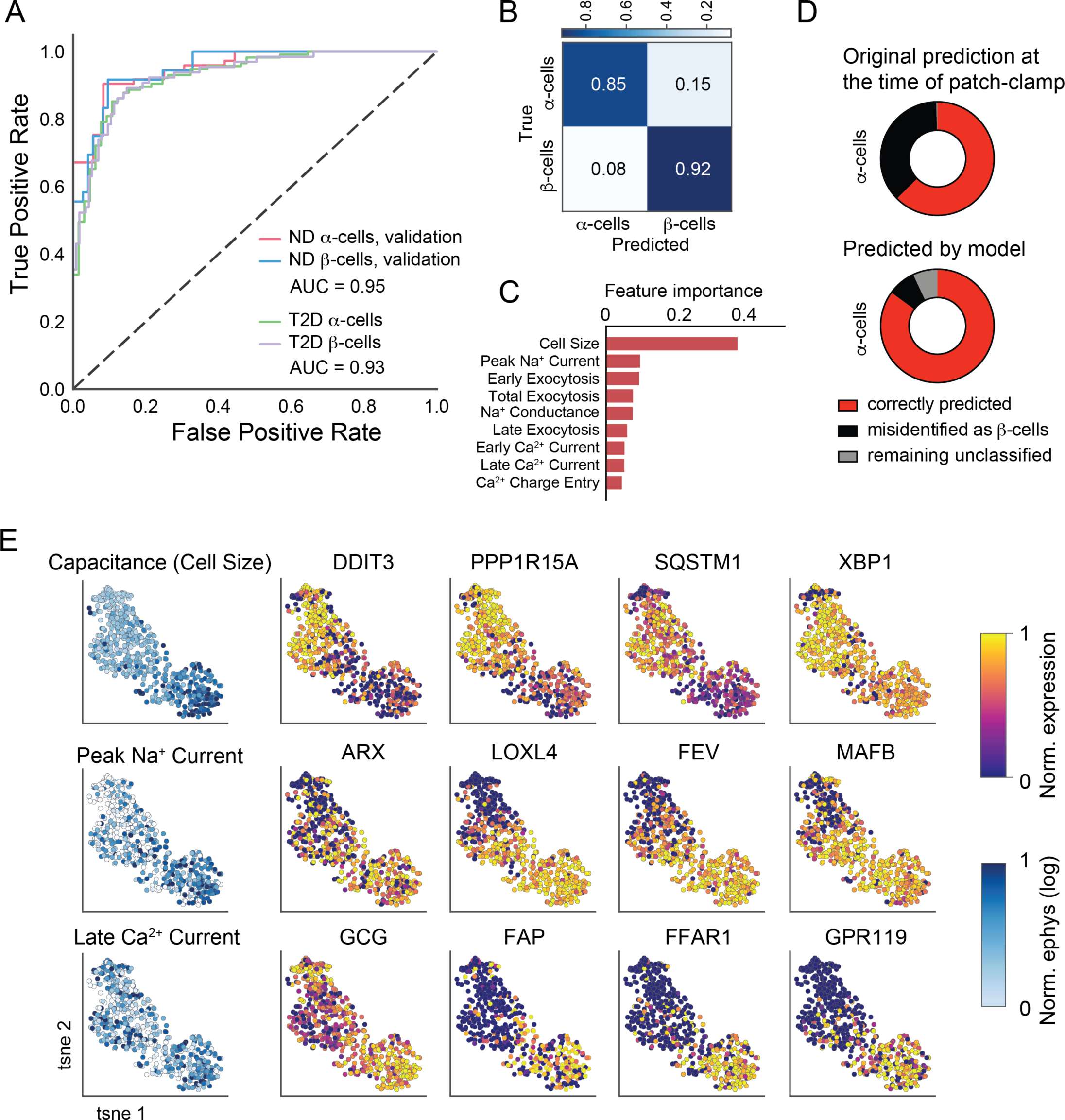
Transcriptomic and electrophysiological heterogeneity in α-cells. (A) ROC curve of cell type prediction using random forests for the validation dataset of ND cells (red, blue) and in T2D cells (green, purple). (B) Confusion matrix in the ND validation dataset. (C) Contribution of each feature to the random forest model. (D) Comparison of α-cell identification from predictions at the time of patch-clamping (simple cell size cut-off) versus the model. (E) t-SNE plots showing heterogeneity in gene expression of α-cells using over-dispersed genes and normalized electrophysiological measurements.

### Application of patch-seq to rare cryo-preserved samples: Islet cells in T1D

Given that human islets retain electrophysiological and transcriptomic integrity following cryopreservation (Manning Fox et al., 2015), we investigated whether the patch-seq approach could be extended to rare tissue types that may only be accessible via tissue banking programs. Single-cell transcriptomes from cryo-banked islets had comparable quality metrics to those of fresh tissue (**Supp Fig 12**), allowing us to investigate cells from three T1D donors and three controls matched for BMI, age, sex and storage-time (**Supp Table S6**). We performed patch-seq in 348 cells from these samples (**Fig 6A; Supp Table S6**) and used a logistical regression model to identify marker genes for each population (**Fig 6B**). Samples from T1D donors contained an unanticipated variety of cell types (**Fig 6B**), including an enrichment for pancreatic polypeptide-secreting γ-cells and ductal cells (**Supp Fig 12**).

**Figure 6.**
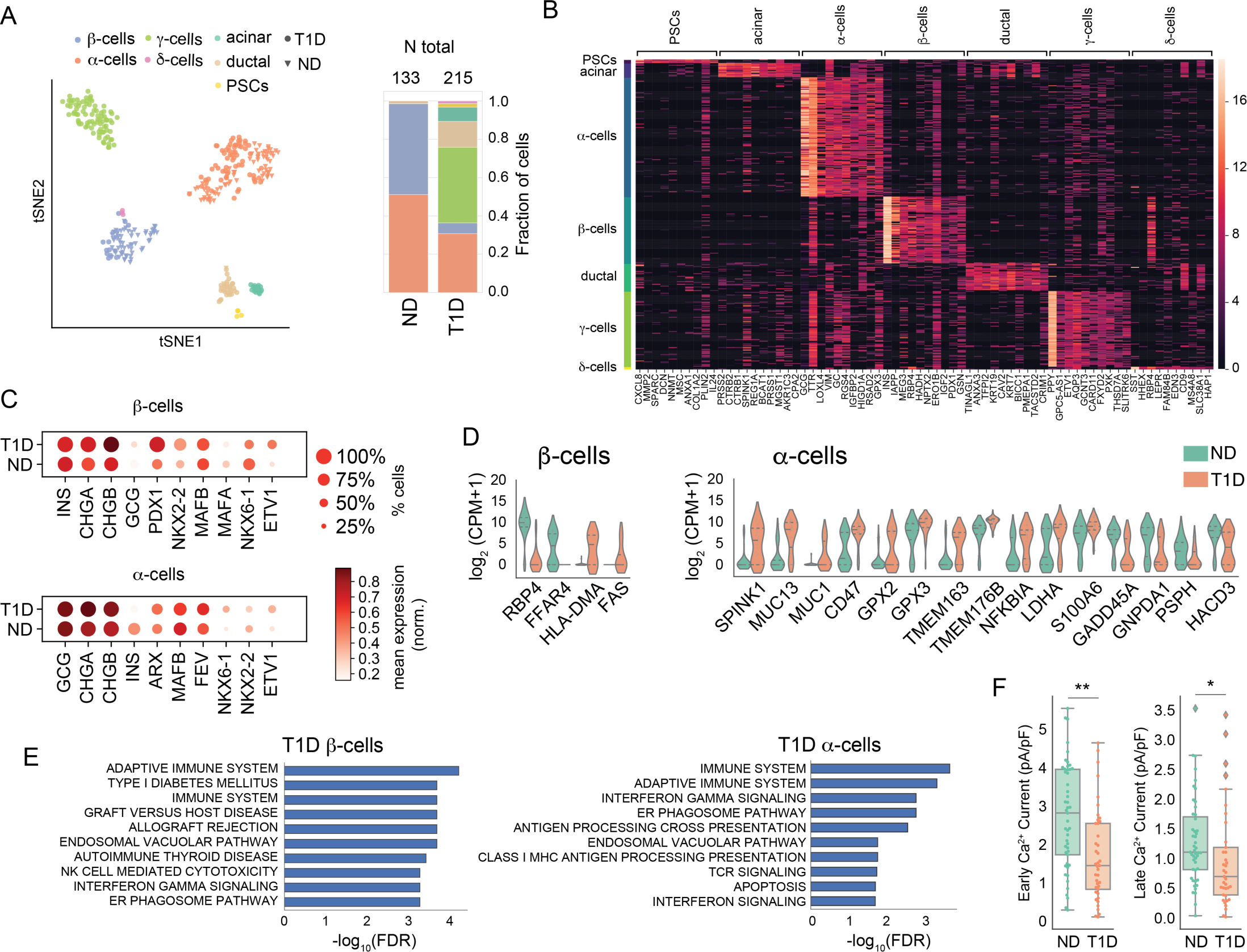
Pancreas patch-seq in cells from cryopreserved T1D islets. (A) *Left:* tSNE projection of measured patch-seq cells clustered by gene expression of over-dispersed genes. *Right:* Distribution of cell types and total number of cells obtained for ND and T1D. (B) Marker genes for each cluster obtained from a logistic regression model. (C) Expression of key identity genes on α- and β-cells from T1D and ND matched controls. (D) Representative genes obtained in a differential expression analysis between T1D and ND for β- and α-cells respectively. (E) Pathways enriched in upregulated genes in T1D α- and β-cells. (F) Distribution of calcium parameters showing statistically significant differences between α-cells of T1D and ND. ** FDR<0.01, * FDR<0.05. Mann-Whitney-U test with BH correction.

We obtained 11 β-cells from two T1D donors, consistent with prior observations of β-cell survival years after T1D onset (Keenan et al., 2010; Morgan and Richardson, 2018) (**Supp Fig 12**). These β-cells had similar electrophysiological profiles as cryopreserved control ND β-cells (**Supp Fig 12**), and appeared to maintain equivalent expression of hallmark β-cell genes (*INS, PDX1, MAFB*) (**Fig 6C**). Differential expression analysis showed decreased expression of *RBP4* and *FFAR4* and transcript enrichment of genes related to immune activation and allograft rejection (*HLA-DMA*, *FAP*) (**Fig. 6D, 6E**). In T1D α-cells we found increased *NKX6.1* and decreased *NKX2.2* mRNA (**Fig 6C; Supp Fig 12**) along with decreased Ca^2+^ channel activity (**Fig 6F; Supp Fig 12**), in general agreement with recent reports using bulk RNA-seq and immunohistochemistry (Brissova et al., 2018; Chakravarthy et al., 2017), although we did not detect a consistent decrease in Ca^2+^ channel gene expression as previously reported (**Supp. Fig. 12**). Our analysis of T1D α-cells also showed transcript enrichment of mucin (*MUC*) and other genes typically associated with ductal cells, and *FEV1*, a reported endocrine progenitor cell marker in mice (**Fig. 7D**; **Supp Table S7**) (Byrnes et al., 2018; Liu et al., 2018). Together, our data demonstrates the use of patch-seq with rare cryo-stored tissue types, and support the view that the transcriptomic and functional signatures of surviving β-cells may be preserved in T1D, while α-cells may have loss of characteristic functional and transcriptomic phenotypes, consistent with the observation of impaired glucagon regulation in T1D.

## Discussion

Success in identifying the genetic basis of islet function and dysfunction in diseases like diabetes has been limited by an inability to connect islet cell physiologic function with transcriptome regulation. Elucidating mechanisms underlying diabetes, whether common forms like T2D, or rarer forms like T1D, also suffers from limited human cell and tissue availability that hinders single cell-based investigations. Multi-modal single-cell technologies will help address these issues by increasing the depth of available data and by making it possible to directly link transcriptomes to additional read-outs in individual cells (Macaulay et al., 2017; Stuart and Satija, 2019). One integrated approach previously used in studies of rodent brain slices is dual patch-clamp electrophysiology and single-cell sequencing (patch-seq) (Cadwell et al., 2016; Földy et al., 2016; Fuzik et al., 2016). Like neurons, pancreatic islet cell secretion is coupled to dynamic electrophysiological mechanisms, making patch-clamp a versatile tool for islet cell functional assessment. Here, we describe development of pancreas patch-seq, and report paired functional-transcriptomic data for more than 1,300 human islet-cells. This depth of data allowed us to link transcriptional heterogeneity to functional heterogeneity at single-cell resolution, and to identify genes and pathways regulating exocytosis of nondiabetic human islet β and α cells. With a patch seq-derived minimal Predictive Set of genes and machine-based learning tools, we developed algorithms that correctly predicted hallmark physiological functions like exocytosis from gene expression, and accurately classified live isolated islet cell types. For studies of islets from T1D and T2D donors, we used patch-seq for unprecedented analyses that identified molecular mechanisms likely to underlie β-cell and α-cell dysfunction in these diseases. Thus, our study provides a powerful heuristic resource and toolset for simultaneous multiplex phenotyping of human islet cells, in health or disease.

Patch-seq analysis of electrophysiological and transcriptomic measurements in single cells identified key genes and pathways regulating β-cell exocytosis. These included both known and previously uncharacterized regulators and mechanisms controlling insulin secretion, such as cell adhesion molecules and neuroactive ligand receptors (**Fig 2C, Supp. Fig 5, Supp. Table 2**). Adhesion molecules have previously-demonstrated roles in β-cell attachment and motility (Dahl et al., 1996; Kaido et al., 2004), and are thought to govern β-cell polarity and function through control of directed insulin exocytosis into the vascular axis (Geron et al., 2015; Parnaud et al., 2015; Gan et al., 2018). Among neuronal regulators, we identified pairs of factors such as neurexin 1 (*NRXN1*) and neuroligin 1 (*NLGN1*), which are known to bind in a heteromeric complex (Suckow et al., 2008). The identification of binding partners in our gene-function correlation analysis confirms the robustness of patch-seq as a tool to identify unanticipated candidate regulators of islet-cell function. Similarly, examination of tissue-specificity reveals that several of the encoded proteins correlating to β-cell exocytosis are also enriched in other excitable cell-types (*GPRASP1*, *RGS9* and *MRAP2* in retinal tissue, *GPRIN1* in brain, *KCNH2* in cardiomyocytes) (Uhlen et al., 2015), indicating shared mechanisms of signal transduction in multiple excitable cell types, and showing that the analytical tools developed here could be applicable to patch-seq studies in other excitable cells and tissues.

Regulation of islet exocytosis by metabolism is complex (Newgard, 2017). We observed multiple expected metabolic pathways to positively correlate to β-cell functional measurements (e.g., glucose flux and metabolism, circadian rhythm; **Fig 2C, Supp. Fig 6**), but also observed unexpected negative correlations of β-cell exocytosis to pathways like glycolysis and pyruvate metabolism (**Fig 2C, Fig 5E, Supp Fig 9**). We integrated several of the observed metabolic correlations to β-cell exocytosis in a functional map (**Fig 2D**), providing support for the hypothesis that glucose is preferentially used for TCA anaplerosis through pyruvate carboxylase (PC) to facilitate insulin secretion (Alves et al., 2015). These findings are also sustained by measures of *CPT1A* and *SLC16A9* expression (**Supp Table S2**), which mediate the rate-limiting step of fatty-acid oxidation and carnitine efflux, respectively (Aichler et al., 2017; Newgard, 2017). Overall, the observation of glucose homeostasis pathways across our analysis also resonates with the importance of metabolites in shifting transcriptional states of β-cells, recently dubbed “metabolic memory” (Rosen et al., 2018; Sharma and Rando, 2017).

Another useful outcome here is our development of a network analysis tool that links a predictive set (PS) of 484 genes with applied machine-learning techniques. We used this tool to predict β-cell function solely from RNA transcript abundance. Genetic regulators of β-cell activity, including many uncharacterized in islet biology, were identified in the predictive set, and included antioxidant molecules (*GPX3*, *GSTK1*), surface proteins (*TSPAN1*, *TSPAN2*), and β-cell enriched molecules (*NPTX2*, *FAM159B*; **Fig 2E**; **Supp Materials S1; Supp Table S2**). To validate predictions from patch-seq correlations and this network analysis, we used genetic loss-of-function approaches, which provided evidence confirming that genes like *OGDHL, RGS9, TSPAN1* and *FAM159B* are positive regulators of β-cell exocytosis (**Fig 2E**). Prior studies have shown that *FAM159B* may be regulated by the β-cell transcription factor SIX3 (Arda et al., 2016), and is a marker of β-cell subsets (Dorrell et al., 2016). Like *FAM159B*, other genes linked by our analysis to differential β-cell function were also found to be expressed in β-cell sub-populations (*ID2, ID4, RBP4*) (Baron et al., 2016; Dorrell et al., 2016; Rui et al., 2017; Segerstolpe et al., 2016). For *RBP4*, we provide evidence that ND RBP4^−^ β-cells, compared to ND RBP4^+^ β-cells, have relatively superior electrophysiological function (also see discussion of T1D below).

Understanding the basis of islet adaptation or dysfunction in diabetes is another important application of patch-seq described here. With increased metabolic demand in physiological or pathological settings, β-cells can compensate by enhancing their excitability, Ca^2+^ signalling, and exocytosis (Chen et al., 2016). Our observation that transcripts positively-correlated with exocytosis in ND β-cells are induced in β-cells of subjects with recent onset T2D, and that transcripts negatively-correlated to ND β-cell exocytosis are reduced in T2D (**Fig 4C**), suggests molecular mechanisms for functional compensation by β-cells in early-stage T2D. However, our analysis also reveals as-yet unexplained complexity in β-cell responses; for example, *ETV1* and *STAT3* mRNA are positively-correlated with ND β-cell functional phenotypes, and also with T2D β-cell dysfunction. Recent studies suggest that β-cell ETV1 and STAT3 protein degradation are regulated by the ubiquitin ligase complex (Dallavalle et al., 2016; Suriben et al., 2015). Our work suggests that ubiquitination and proteosomal degradation pathways in β-cells, including a pathway stimulating COP1 auto-degradation, are associated with dysfunction in T2D (**Fig 4**; **Supp Fig 10**). Thus, in addition to transcript-based mechanisms, our results suggest that post-translational mechanisms involving factors like COP1, ETV1 and STAT3 may govern β-cell dysfunction or responses in T2D (O’Shea and Plenge, 2012; Saarimäki-Vire et al., 2017; Suriben et al., 2015).

Patch-seq also advances our understanding and ability to study human α-cells. Cell size and Na^+^ channel properties are key identifiers used to distinguish rodent α- and β-cells (Briant et al., 2017; Zhang et al., 2014), but human α-cell heterogeneity results in overlap of these electrophysiological properties (**Fig 1F**). Using patch-seq data, we developed additional models for improved cell-type identification of live human islet α-cells and β-cells (**Fig 5A-D**). We also show that specific phenotypes like Na^+^ current activities and cell size (**Fig 5E**) can vary significantly in α-cells, and that this functional heterogeneity corresponds well with expression of transcripts specifying islet cell lineage (*ARX, MAFB, FEV*) or governing cellular stress (*DDIT3, PPP1R15A*). Prior studies have reported the impact of the ER-stress response in β-cells subpopulations (Baron et al., 2016; Muraro et al., 2016), and in α-cells (Akiyama et al., 2013; Burcelin et al., 2008; Korsunsky et al., 2018), suggesting that varying levels of ER-stress could drive altered or dysfunctional states in both islet cells types. We noted α-cell subpopulations expressing markers indicating either low- or high ER-stress, both in ND and T2D donors. Thus, patch-seq merged electrophysiological and transcriptomic data to document, and suggest a basis for, heterogeneous function and transcriptome regulation in human α-cells.

Phenotyping live islet cells from donors with T1D and other forms of diabetes is a significant challenge, stemming from the rarity of available tissue, and low islet cell recovery in those settings; this has retarded understanding of islet and diabetes biology (Brissova et al., 2018; Wang et al., 2016b). Likewise, procuring and studying appropriate control tissue for studies of T1D islets, matched in key properties like donor age, is an enduring challenge. Electrophysiological studies of islets from only one T1D donor have been reported, suggesting that surviving β-cells have normal function and α-cells are dysfunctional (Walker et al., 2011). Similarly, to date only islets from one donor have been studied using single-cell RNAseq (Wang et al., 2016b). Thus, our extension of patch-seq to studies of cryo-stored islets from ND and T1D donors (Manning Fox et al., 2015) represents a significant innovation. Here we show that library quality from cryo-preserved islets is comparable to that of control fresh islet cells (**Supp Fig 12**). Moreover, our detection of α-cell dysfunction and normal function of surviving β-cells using patch-seq with T1D islets correlates well with findings from a recent study using different methods (Brissova et al., 2018). Patch-seq revealed greater α-cell heterogeneity in T1D compared to ND islets, including evidence of impaired maintenance of α-cell fate; that might contribute to the well-recognized impairment of glucagon secretion observed in T1D (Unger and Cherrington, 2012). Moreover, transcriptomic profiling here reveals that functional preservation in T1D β-cells corresponds to undetectable expression of *RBP4* (**Fig 6**), consistent with work here and by others (Brissova et al., 2018; Rui et al., 2017). By contrast, we show that T1D ductal cells, which some suggest represent a potential source of new β-cells (Corritore et al., 2016), lack detectable endocrine physiological phenotypes (**Supp Fig 12**). Whether from pancreatic cell reprogramming or other sources (Chakravarthy et al., 2017; Thorel et al., 2010), rigorous evaluation of “replacement” β-cells with patch-seq should emerge as a new benchmark to assess functional and transcriptional resemblance to *bona fide β*-cells.

In conclusion, the patch-seq approach for pancreas integrates multiple assays of islet-cells in health and diabetes. Thus, our work represents an important step in the development of multimodal scRNAseq technologies, by increasing cell throughput and extending analyses of simultaneously-measured transcriptomic data and physiological features. The approaches and data presented here help clarify observed inherent islet cellular heterogeneity, and provide valuable tools for understanding the molecular mechanisms underlying normal islet function and dysfunction in diabetes at single-cell resolution.

## Methods

### Islet isolation, cryopreservation, and insulin secretion

Donor organs were obtained with written consent and research ethics approval at the University of Alberta (Pro00013094, Pro00001754), perfused via the pancreatic ductal system with buffer containing Collagenase Gold 800 (VitaCyte, Indianapolis, IN) and Thermolysin (Roche Diagnostics, Mannheim, Germany), then digested with a Ricordi Islet Isolator (Biorep Diabetes, Miami, FL) and purified by density centrifugation. Donor characteristics are described in **Supp Tables S1,S3,S6**. Full details of our human islet isolation protocol, equipment setup, quality control, cryopreservation, and static glucose-stimulated insulin secretion assays have been deposited in the protocols.io repository (Lyon et al.).

### Electrophysiological phenotyping

Hand-picked human pancreatic islets were dissociated to single cells using enzyme-free Hanks’-based Cell Dissociation Buffer (Thermo Fisher Scientific, Cat#13150-016, for donors R229~R242) or Hanks’ Balanced Salt Solution and StemPro accutase (Thermo Fisher Scientific, Cat#A11105-01, for donors R243~R269, cryo-T1D donors and matched controls) and cultured in low glucose (5.5 mmol/L) DMEM with L-glutamine, 110 mg/L sodium pyruvate, 10% FBS, and 100 U/mL penicillin/ streptomycin for 1-3 days. Media were then changed to bath solution containing (in mM): 118 NaCl, 20 TEA, 5.6 KCl, 1.2 MgCl_2_•6H_2_O, 2.6 CaCl_2_, 5 HEPES, and either 1, 5 or 10 glucose (pH 7.4 with NaOH) in a heated chamber (32–35°C). For whole-cell patch-clamping, fire polished thin wall borosilicate pipettes coated with Sylgard (3-5 MOhm) contained intracellular solution with (in mM): 125 Cs-glutamate, 10 CsCl, 10 NaCl, 1 MgCl_2_•6H_2_O, 0.05 EGTA, 5 HEPES, 0.1 cAMP, and 3 MgATP (pH 7.15 with CsOH). Electrophysiological measures were collected using a HEKA EPC10 amplifier and PatchMaster Software (HEKA Instruments Inc, Lambrecht/Pfalz, Germany) and protocols shown in **Fig 1E** within 5 minutes of break-in. Quality control was assessed by the stability of seal (>10 GOhm) and access resistance (<15 MOhm) over the course of the experiment. Data were analysed using FitMaster (HEKA Instruments Inc) and Prism 6.0h (GraphPad Software Inc., San Diego, CA). Immediately following recordings, the patch pipette was withdrawn and replaced with a wide-bore (0.2-0.5 MOhm) collecting pipette containing lysis buffer without ERCC mix. Cells were then collected by gentle suction and visual confirmation, and then transferred into 8-strip PCR tubes containing 4 μl lysis buffer (with ERRC spike-in) on ice and stored at −80°C until cDNA and library preparation (see below).

### siRNA transfection

Dissociated human islets cells were transfected with scramble siRNA (Cat# 1027284, Qiagen, Toronto, Canada) or siRNA from ThermoFisher Scientific against OGDHL (ID# s31422), TSPAN1 (ID# s19659), FAM159B (ID# 264018), RGS9 (ID# s16736), and GYG1 (ID# s6360) together with a fluorescent marker (Allstars Neg.siRNA AF488, Qiagen, Cat# 1027292) using DharmaFECT 1 (GE Healthcare) according to manufacturer’s protocol. In patch-clamp studies, the visible fluorescence was checked to identify positively transfected cells. Total RNA was prepared by TRIzol (Invitrogen) according to manufacturer’s protocol. The cDNA was prepared from 100-200 ng of total RNA using 5xAll-In-One RT Master mix (Applied Biological Materials Inc). Real-time PCR was performed using Fast SYBR Green Master Mix, 7900HT Fast Real-Time PCR systems (Applied Biosystems) and primers designed to flank an intron of each gene to confirm knockdown (**Supp Table S4**).

### Islet dispersion for FACS and single-cell library preparation

Human islets were washed once in cold PBS and dissociated into single cells by enzymatic digestion with Accumax (Invitrogen), followed by digestion using freshly prepared Dispase (Fisher Scientific). Cells were filtered using a 70 μm cell strainer, quantified, and stained with LIVE/DEAD Fixable near-IR dead cell dye (Life Technologies, L10119) as a viability marker. Cells were then blocked with mouse IgG in FACS buffer (2% FBS, 10mM EGTA, in PBS), followed by staining with appropriate antibodies at 1:100 (v/v) final concentration. The following combination of antibodies was used to select endocrine cells: HPi2-Alexa-405 (Novus, NBP1-18946AF405), HPx1-Alexa-647 (Novus, NBP1-18951AF647), CD133/1-APC (Miltenyl Biotec, 130-113-668), CD133/2-APC (Miltenyl Biotec, 130-098-129), CD31-APC-Cy7 (BioLegend, 303119). Cells were then sorted using a Sony SH800 cell-sorter and a 100 μm nozzle following doublet removal. Single cells were sorted directly into 384-well plates (Bio-Rad HSP3841) containing 0.4 μL of lysis buffer with dNTPs (Invitrogen) and ERCC spike-in control (ThermoFisher). Plates were centrifuged and placed on dry ice immediately before storage at −80°C.

For both patch-seq cells and FACS-collected cells, we generated cDNA and sequencing libraries using the SmartSeq-2 protocol as previously described (Picelli et al., 2014). For patch-seq cells, we first assembled the 8-strip tubes into 96-well plates for increased throughput (Bio-Rad, RC9601 and MSA5001). For FACS-collected cells we proceeded directly with the obtained 96-well or 384-well plates. Briefly, mRNAs were primed with an anchored oligo-dT and reverse transcribed using an LNA-containing template switching oligo, followed by PCR amplification (21 cycles). Libraries were then generated from the amplified cDNA by tagmentation with Tn5. Libraries were sequenced either in a NextSeq 500 or NovaSeq platform (Illumina) using paired-end reads (75 bp) to an average depth of 1 million reads per cell.

### Processing, quality control and filtering of single-cell RNA-seq data

Sequencing reads were aligned to the human genome (GRCh38 genome with supplementary ERCC sequences) using STAR (Dobin et al., 2013), and gene counts determined using htseq-count (intersection-nonempty) using a GTF annotation with Ensembl 89 release genes (Anders et al., 2015). Analysis of splicing was performed based on the splice junctions called by STAR. Gene expression was normalized to counts per million (cpm) after removal of counts corresponding to ERCC spike-ins, and transformed to log2 values after addition of a pseudocount.

We filtered patch-clamped cells based on stringent QC criteria: >1,500 genes, >100,000 human mapped reads, >40% uniquely mapped reads (STAR), <40% unmapped reads (STAR), and more than 25% of total reads mapped to exons (ht-seq). This filtering retained 89% of all patch-clamped cells (1126/1275). We also filtered cells with low expression of ACTB and GAPDH (<3SD below mean) (29 cells), and potential doublets determined from high levels of hormone coexpression (log_2_CPM>13 for more than one marker gene: INS, GCG, PPY, SST, PRSS1) (76 cells). In this way, we obtained 1,021 high-quality patch-seq transcriptomes for further analysis. For cryopreserved cells, we applied the same QC filtering and obtained equivalent percentages in each step. This provided an additional set of 348 cryopreserved patch-seq cells (411 initial cells).

### Clustering and cell type determination

Initial clustering of cell types was performed by selection of over-dispersed genes (top 2000 genes based on coefficient of variation), followed by dimensionality reduction by PCA (10 PCs), and tSNE projection (perplexity=30, learning rate=200). Clusters were selected based on the Louvain algorithm for community detection or hdbscan, and cell types assigned based on the expression of key marker genes. Selection of marker genes for the T1D clusters was performed using a logistic regression model (python package scanpy) (Wolf et al., 2018). Downstream analysis was performed using custom python and R scripts.

### Correlation between electrophysiological parameters and gene expression

The relationship between electrophysiological parameters and gene expression was measured using rank correlation statistics. Spearman’s correlations were computed in different groups of cells according to cell type, disease condition and experimental protocol (e.g. β, ND, high-glucose condition). We first removed low expressed genes by selecting genes with mean expression log_2_(CPM)>1, corresponding to a total of 3000-8000 genes for dataset. Outliers in electrophysiology were removed by quantile filtering of the highest and lowest 3% of cells for each parameter. We observed outlier cells to be consistent across sets of co-measured parameters (exocytosis, Ca^2+^, Na^+^ currents) suggesting that they might reflect technical noise. Measurements of exocytosis and ionic currents were normalized to initial cell capacitance to account for effects of cell size. For exocytosis measurements, we also collapsed negative values to zero to reduce their effect on the measured correlations, as variations around zero are unlikely to be driving functional responses (i.e. exocytosis). Finally, all current measurements (Ca^2+^, Na^+^) were transformed to positive defined values -regardless of their flow direction across the cell membrane, so positive correlations are representative of genes driving larger responses in electrophysiology.

Spearman tie-corrected correlations were computed for each gene and significance was tested by bootstrapping (1,000 iterations). Reported values are mean, standard deviation, and equivalent z-score from the bootstrapped values. To verify that correlations determined on this way report genes that show significant variation across each electrophysiological parameter, we recomputed correlations after performing a median smoothing of the data. In this case, we sorted all cells according to each electrophysiological parameter and performed a median average with a rolling-window corresponding to 10% of the cells. We then computed correlations and confidence intervals using the same bootstrap approach, finding overlapping gene lists and significance values. We also verified that a bootstrap of donors (instead of cells), provided an overlapping list of genes for total exocytosis in ND β-cells. To further refine the final list of genes for knock-down validation we focused on genes observed in >30% of cells. Analysis was implemented in python scripts.

### Electrophysiological predictions using PS gene set

For patch-seq electrophysiological prediction, we split ND β-cells into a training set (80%) and test set (20%). We used the training set to perform the bootstrapped correlation analysis between electrophysiological parameters and gene expression as detailed above. We selected genes showing significant correlations (|z-score|>2) with at least two electrophysiological parameters belonging to different functional groups, and retained genes with highest median expression (observed in >50% of cells). In this way, we obtained our final PS gene set (484 genes, **Supp Table S5**). We then built a k-nearest neighbors (*k*-NN) model (Pedregosa et al., 2011), to determine the *k* closest cells in PS gene expression to each cell (*k*=5, metric spearman correlation) using the training set. We used the k-NN model to infer the electrophysiological parameters of each cell from the averaged values of the identified neighbors (after masking cells for which an electrophysiological parameter could not be measured). The test set was used for final validation by predicting their electrophysiological parameters using the k-NN model build with the training dataset.

### Statistical analysis

Differential expression was performed using a non-parametric Mann-Whitney U test. For comparisons involving few donors or showing bias in sex representation (e.g. T1D α-cells), we repeated the analysis with VOOM-LIMMA using sex as a covariate. All p-values were corrected for multiple hypothesis testing using Benjami-Hochberg (BH) and significance was determined from a permutation test by shuffling the labels and using an FDR<0.05. Statistical differences between electrophysiological parameters was determined using a non-parametric Mann-Whitney U and corrected for multiple hypothesis testing using BH across all measured parameters unless otherwise stated.

## Acknowledgments

This work was supported by the funding from the Chan Zuckerberg Biohub and the California Institute for Regenerative Medicine to SRQ; by a Foundation Grant from the Canadian Institutes of Health Research (CIHR: 148451) to PEM; and by grants from the U.S. National Institutes of Health (1U01DK10830001 1R01DK107507, 1R01DK108817 and P30 DK116074) to SKK, (1U01DK120447-01) to SKK, SRQ, and PEM, and from JDRF (2-SRA-2019-698-S-B) to SKK and PEM. Support from the Human Islet Research Core in the Stanford Diabetes Research Center is gratefully acknowledged. Human islet isolation at the Alberta Diabetes Institute IsletCore was subsidized by the Alberta Diabetes Foundation.

We thank Dylan Hendersson (Chan Zuckerberg Biohub) for help in sample processing, and Dr. Norma Neff (Chan Zuckerberg Biohub) and Jennifer Okamoto (Chan Zuckerberg Biohub) for sequencing expertise. We thank Dr. Linford Briant (University of Oxford) for helpful discussions on islet cell type modelling; and Dr. Jocelyn Manning Fox (University of Alberta) for critical reading of the manuscript. We thank the organ procurement organizations across Canada, particularly the Human Organ Procurement and Exchange (HOPE) program in Edmonton and the Trillium Gift of Life Network (TGLN) in Ontario, for their work in obtaining human pancreas for research. We also thank Dr. Manning Fox and Mrs. Nancy Smith (University of Alberta) for their contributions to human islet isolations. Finally, we are indebted to organ donors and their families for their generous support of scientific research.

## Data availability

The datasets generated and analyzed in the study are available in the NCBI Gene Expression Omnibus (GEO) and Sequence Read Archive (SRA) and can be accessed upon request.

## Author Contributions

JCS and XQD developed the patch-seq pipeline, XQD performed and analyzed patch-clamp data, JCS performed libraries and analyzed patch-seq data, XQD and KS performed knockdown experiments, JCS and YH performed FACS, JL and AB performed islet isolations, AB performed insulin measurements, JL performed cryopreservation, JCS, XQD, YH, SK, SRQ and PEM discussed data and interpreted results, SK, SRQ and PEM supervised the project, and all authors wrote the manuscript.

## Author Disclosures

The authors disclose no conflict of interest.

