## Supplemental Information and Figures for "Pancreas patch-seq links physiologic dysfunction in diabetes to single-cell transcriptomic phenotypes"

### Supplementary Table S1. Donors included in initial patch-seq experiments.

| DonorID | Diabetes | Age (years) | Sex | BMI | HbA1c | Time from diagnosis (years) | Treatment (if known) | Donation Type | Patched cells | FACS cells | Reason for exclusion (if applicable) |
|---|---|---|---|---|---|---|---|---|---|---|---|
| R229 | NO | 22 | F | 23.0 | 5.3 | n/a | n/a | NDD | 18 | NO | |
| R230 | NO | 58 | M | 29.4 | 6.2 | n/a | n/a | NDD | 58 | YES | |
| R232 | NO | 66 | F | 18.5 | 6.1 | n/a | n/a | NDD | 66 | NO | |
| R233 | NO | 76 | F | 19.3 | 6 | n/a | n/a | NDD | 76 | NO | |
| R234 | NO | 50 | F | 31.7 | 5.7 | n/a | n/a | NDD | 35 | NO | |
| R235 | NO | 53 | F | 24.5 | 5.7 | n/a | n/a | NDD | 68 | YES | |
| R237 | NO | 61 | M | 19.7 | 5.9 | n/a | n/a | NDD | 54 | NO | |
| R238 | NO | 52 | M | 26.1 | 5.7 | n/a | n/a | NDD | 23 | NO | |
| R239 | NO | 24 | F | 22.0 | 5.5 | n/a | n/a | NDD | 46 | NO | |
| R242 | NO | 46 | M | 20.1 | 5.9 | n/a | n/a | DCD | 16 | YES | |
| R243 | NO | 39 | M | 27.1 | 5.8 | n/a | n/a | DCD | 40 | NO | |
| R246 | NO | 65 | F | 39.2 | 5.8 | n/a | n/a | NDD | 93 | NO | |
| R247 | NO | 72 | M | 23.9 | n/a | n/a | n/a | NDD | 23 | YES | |
| R252 | NO | 26 | F | 25.4 | 5 | n/a | n/a | NDD | 54 | YES | |
| R253 | NO | 57 | M | 25.6 | 5 | n/a | n/a | NDD | 64 | YES | |
| R256 | NO | 23 | M | 32.5 | 5.4 | n/a | n/a | NDD | 49 | YES | |
| R260 | NO | 73 | F | 26.9 | 6.2 | n/a | n/a | NDD | 44 | NO | |
| R264 | NO | 44 | M | 33.8 | 5.7 | n/a | n/a | NDD | 56 | YES | |
| R231 | T2D | 41 | F | 37.1 | 6.8 | 0.5 | diet only | NDD | 35 | YES | |
| R240 | T2D | 55 | F | 30.9 | 6.9 | 2 | diet only | NDD | 90 | NO | |
| R241 | T2D | 65 | M | 21.8 | 9.9 | 25 | insulin | NDD | 21 | YES | |
| R244 | T2D | 48 | F | 30.5 | 7.5 | 3 | metformin | NDD | 71 | YES | |
| R257 | T2D | 56 | M | 30.2 | 7.3 | undiagnosed | untreated | DCD | 35 | YES | |
| R259 | T2D | 45 | F | 33.1 | 6.4 | 4 | oral medications | NDD | 38 | YES | |
| R263 | T2D | 71 | M | 38.3 | 7.6 | 9 | metformin | NDD | 21 | YES | |
| R262 | other | 54 | F | 22.4 | 5.6 | n/a | | NDD | 93 | NO | Insulin-treated T2D lost significant weight and reverted to metformin-only |
| R265 | other | 64 | M | 37.7 | 6.6 | n/a | | DCD | 86 | YES | Parkinson’s disease |
| R269 | other | 14 | M | 21.5 | n/a | n/a | | DCD | 78 | NO | Age |

***NDD – Neurological Determination of Death; DCD – Donation after Cardiac Death*

### Supplementary Table S3. Donors used in siRNA validation studies.

| DonorID | Diabetes | Age (years) | Sex | BMI | HbA1c | Donation Type | siRNA experiments used for |
|---|---|---|---|---|---|---|---|
| R278 | NO | 57 | M | 27.6 | 5.7 | NDD | siOGDHL |
| R281 | NO | 30 | F | 25.1 | n/a | NDD | siOGDHL |
| R282 | NO | 57 | M | 26.4 | 6.0 | NDD | siFAM159B; siRGS9; siGYG1 |
| R283 | NO | 22 | M | 22.5 | n/a | NDD | siTSPAN1; siFAM159B; siRGS9 |
| R285 | NO | 19 | M | 21.6 | 5.4 | NDD | siTSPAN1; siFAM159B; siRGS9; siGYG1 |
| R286 | NO | 41 | M | 20.4 | 5.2 | NDD | siTSPAN1; siGYG1 |
| H2182 | NO | 48 | F | 38.5 | n/a | NDD | siOGDHL |
| H2191 | NO | 44 | M | 21.1 | n/a | NDD | siTSPAN1; siFAM159B; siRGS9; siGYG1 |

**NDD – Neurological Determination of Death; DCD – Donation after Cardiac Death*
**donor beginning with HXXX is from the clinical islet laboratory at the University of Alberta*

### Supplementary Table S4. Primers used for quantitative PCR.

| Gene Name | Accession | 5'-3' primer sequence | Amplicon length (bp) | % knockdown (avg+/-STE) |
|---|---|---|---|---|
| OGDHL | NM_018245.2 | FW - GATGTGGAGCAGTGCCAGTG<br>RV - ATGGAGTGGCAGCTAGGCAC | 129 | 78.5+/-3.1 (n=3) |
| TSPAN1 | NM_005727.3 | FW - GTGTACACCACAATGGCTGAGCACTTC<br>RV - CGTATAGTTGGTGAAGCCACAGCACTTGAG | 144 | 87.4+/-0.4 (n=4) |
| FAM159B | NM_001164442.1 | FW - CCTACAAGCACAGCTACATGTGGAGC<br>RV - GGACACAGACACTGATGACAAAGGCGAG | 102 | 78.4+/-11.1 (n=4) |
| RGS9 | NM_003835.3 | FW - CTGGACCGAGTGACCAATCCGAATG<br>RV - CACTGTGGACCTCATCAAGGCCT | 111 | 86.4+/-3.2 (n=3) |
| GYG1 | NM_004130.3 | FW - TCCACTGCTGGTCGCTTACACAG<br>RV - GGTCTGGTGCTGCTGACAATTCTTCTC | 117 | 81.4+/-3.1 (n=4) |

### Supplementary Table S6. Donors included in study of cryopreserved samples.

| DonorID | Diabetes | Age (years) | Sex | BMI | HbA1c | Time from diagnosis (years) | Donation Type | Duration of Cryopreservation (years) | FACS cells |
|---|---|---|---|---|---|---|---|---|---|
| R134 | NO | 23 | M | 21.9 | 5.9 | n/a | DCD | 3 | NO |
| R124 | NO | 43 | F | 24.6 | 5.2 | n/a | NDD | 3 | NO |
| R177 | NO | 28 | M | 23.4 | n/a | n/a | NDD | 2 | NO |
| R119 | T1D | 27 | M | 18.5 | n/a | 11 | NDD | 3 | NO |
| R132 | T1D | 39 | F | 24.6 | 5.9 | 31 | NDD | 3 | NO |
| R079 | T1D | 32 | M | 21.9 | 9.3 | 17 | NDD | 4 | YES |

**NDD – Neurological Determination of Death; DCD – Donation after Cardiac Death*

### SUPPLEMENTARY FIGURE CAPTIONS

**Supp. Fig. 1 – Pancreas patch-seq pipeline.** (A) Scheme of the pancreas patch-seq pipeline to obtain cells from the same donors in 96-well (patch-seq) and 384-well (FACS) formats. (B) Fragment analyzer image showing size distribution of cDNA obtained from patch-seq cells after library preparation (SmartSeq2). (C) Violin plot showing sequencing depth of patch-seq and FACS cells obtained in this dataset (~1 million reads/cell).

**Supp. Fig. 2 – Distribution of measured electrophysiologic parameters across cell types and disease.** Boxplots showing distribution of measured parameters for every cell type: $\alpha$-cells (orange), $\beta$-cells (blue), $\gamma$-cells (green), $\delta$-cells (brown) and acinar cells (pink), separated by disease condition (ND or T2D). Results are aggregated across different glucose conditions, and diamonds denote outliers.

**Supp. Fig. 3 – Quality control metrics and comparison to previous FACS datasets.** (A) Sequencing metrics of patch-seq cells versus FACS controls. Patch-seq cells show comparable % of reads uniquely mapped to the human genome and exonic regions, and do not show an increase of unmappable short reads characteristic of degraded RNA and low-quality libraries. Variation in percentage of ERCC and multimapped reads (ribosomal RNA), are likely due to the 10-fold decrease in volume used in 384-well plates (FACS) versus 96-well plates (patch-seq) (i.e. lower amount of ERCCs in 384-well plates and different priming efficiencies). (B) Number of genes per cell (expression > 3 counts per million) in patch-clamp cells (pink), FACS cells (green), and two previous FACS datasets obtained using Smartseq2 (blue, purple) (Enge et al., 2017; Segerstolpe et al., 2016). (C) Distribution of gene expression of key genes for the main islet cell-types identified: $\alpha$- , $\beta$-, $\delta$-, and $\gamma$-cells (top to bottom panels respectively). For each gene, the distribution of gene expression in the patch-clamp dataset (pink) is compared to FACS cells (green) and two previous FACS datasets Segerstolpe (blue), Enge (purple).

**Supp. Fig. 4 – Effects of single-cell dispersion and patch-clamp on gene expression.** (A,B) tSNE and violin plots of genes showing changes in expression immediately after dispersion (day 0) and after recovery (days 1-3) for $\alpha$- and $\beta$-cells in the patch-clamp dataset. As described in previous single-cell studies (van den Brink et al., 2017), single-cell dispersion induces an increase on immediate early genes (FOSB, JUND), that is reduced after recovery in culture. Genes that show increased expression after recovery include ferritin subunits, ID genes, and islet-identity genes like MAFB. (C) Correlation plot of top over-dispersed genes in donor pseudo-bulk averages (aggregated counts) for all $\alpha$- and $\beta$-cells aggregated by protocol (FACS, patch-clamp) and donor. The plot shows 4 clusters of genes that correspond to each cell-type ($\alpha$-, $\beta$-) or protocol (patch-clamp, FACS) as shown in D. (D) Donor averages of characteristic genes obtained from the four clusters in panel C. Genes that distinguish patch-clamp and FACS (genes highlighted in blue and purple) overlap with those observed in panels A,B, suggesting that most differences are accounted by an increase in IEG after single-cell dispersion. (E) tSNE showing patch-seq cells (light colors) together with controls that underwent the same culture and collection process but that were directly collected without performing patch-clamp measurements (dark colors). Non-patched controls overlap with the overall distribution of patched cells, showing that there are no significant gene expression differences due to patching.

**Supp. Fig. 5 – $\beta$-cell correlation analysis and gene pathways.** (A) Correlation plot of measured electrophysiological parameters in ND $\beta$-cells including donor metadata (Age, HbA1c, BMI, Sex), and experimental conditions in patch-clamp measurements. The measured parameters show consistent clusters for each functional group, and do not show significant cross-correlation to other donor or experimental parameters that could account for confounding effects. (B) Chord plot of pathways correlated to exocytosis parameters (early, late, total, and normalized to $Ca^{2+}$ charge) in ND $\beta$-cells. Each enriched pathway is connected to the genes for which we obtained positive correlation to exocytosis (i.e genes enriched in cells with high exocytosis). Genes are ordered from top to bottom according to correlation significance (z-score).

**Supp. Fig. 6 - Summarized GSEA for each functional group.** GSEA using averaged correlation z-scores for each group as enrichment metric. Pathways with top normalized enrichment are shown (i.e. pathways enriched in positively correlated genes). Inset is FDR from a permutation test (cut-off FDR<0.2).

**Supp. Fig. 7 – Prediction of β-cell electrophysiology from nearest neighbours in gene expression.** Scatter plots for each electrophysiological parameter showing measured (x-axis) versus predicted value (y-axis) from its nearest-neighbours in gene expression (k-NN, n=5). Predictions are first done in a subset of cells (black) for which the 'PS gene set' is obtained and used to calculate the nearest neighbours (484 genes, see Methods). Spearman correlation and p-value for each predicted parameter is indicated in black. Performance is further tested in a reduced set of cells withheld from the previous gene selection and prediction process (red). For this set we obtained significant correlations (p<0.05) for a subset of parameters (total and late exocytosis, late $Ca^{2+}$ conductance, $Na^{+}$ conductance and peak $Na^{+}$ current). Similar p-values are obtained by comparison to a random set of genes of similar expression to the 'PS gene set'.

**Supp. Fig. 8 – Expression of RBP4 in β-cells correlates to decreased $Na^{+}$, $K^{+}$ channel expression and function.** (A) Boxplots of electrophysiological function in beta cells according to expression of the heterogeneity marker RBP4. Individual cells are shown in gray. A significant decrease of $Na^{+}$ currents and exocytosis is observed for RBP4+ cells. (B) Gene expression distribution (top) and % of cells expressing the gene above detection threshold (bottom) for all $Na^{+}$, ATP-sensitive $K^{+}$ channels and $Ca^{2+}$ channels detected in beta cells. We observe a significant decrease of SCN3A, KCNJ8 and ABCC9 in RBP4+ cells. All comparisons are based on a Mann-Whitney-U test with Bonferroni correction for multiple hypothesis testing. *** FDR<0.001, ** FDR<0.01,* FDR<0.05.

**Supp. Fig. 9 – Electrophysiological differences in T2D and correlation analysis.** (A) Boxplot showing distribution of electrophysiological parameters for ND and T2D β-cells. All comparisons are based on a Mann-Whitney-U test with Bonferroni correction for multiple hypothesis testing. *** FDR<0.001, ** FDR<0.01,* FDR<0.05. (B) Genes with reversed correlation (from positive to negative) between ND and T2D for at least one electrophysiological parameter. (C) Genes with concordant positive correlations between ND and T2D for at least one electrophysiological parameter.

**Supp. Fig. 10 – Pathway analysis in T2D and ETV1 expression.** (A) Heatmap showing all enriched pathway for genes that (from left to right): correlate to exocytosis in ND, correlate to exocytosis in T2D, anticorrelate to exocytosis in ND, anticorrelate to exocytosis in T2D. Some immune related pathways are shared between correlated and anticorrelated groups. (B) Distribution of gene expression of ETV1 and fold-change of several genes in dataset from (Segerstolpe et al., 2016). (E) Splicing map of ETV1 in β-cells from our dataset.

**Supp. Fig. 11 – Transcriptomic and physiological heterogeneity in α-cells.** (A) Additional t-SNE plots showing heterogeneous gene expression in α-cells, and additional electrophysiological measurements not show in the main panel. (B) Spearman correlation of measured electrophysiological, donor metadata and experimental conditions across all patch-seq α-cells. All electrophysiological parameters are normalized by cell size.

**Supp. Fig. 12 – Patch-seq on cryopreserved T1D samples.** (A) Sequencing metrics of patch-seq cryopreserved cells versus fresh patch-seq cells. (B) Violin plots for marker genes obtained for each cell type using a logistic regression model. (C) Distribution of cell types and total number of cells obtained for each donor. (D) Boxplots showing distribution of measured electrophysiological parameters for every cell type: α-cells (orange), β-cells (blue), γ-cells (green), ductal cells (brown), δ-cells (pink) and PSCs (yellow), separated by disease condition (ND or T1D). All experiments performed at 5 mM glucose. (E) Enrichment analysis between alpha cells from T1D and ND donors for the genes reported in (Brissova et al., 2018). Data corresponds to the log fold change of pseudo-bulk averages (aggregated counts) for all α-cells.

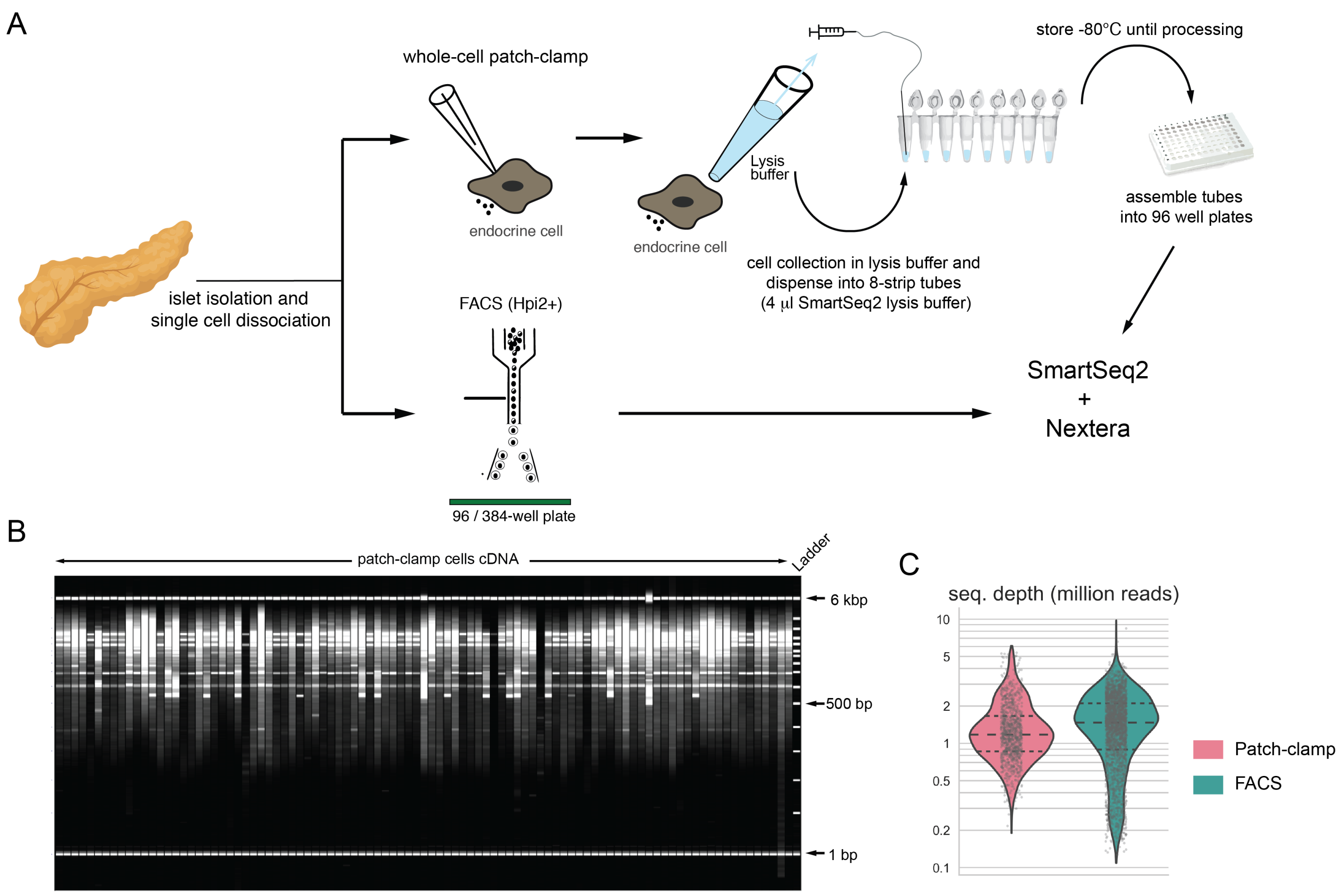


**Supp. Figure 1**

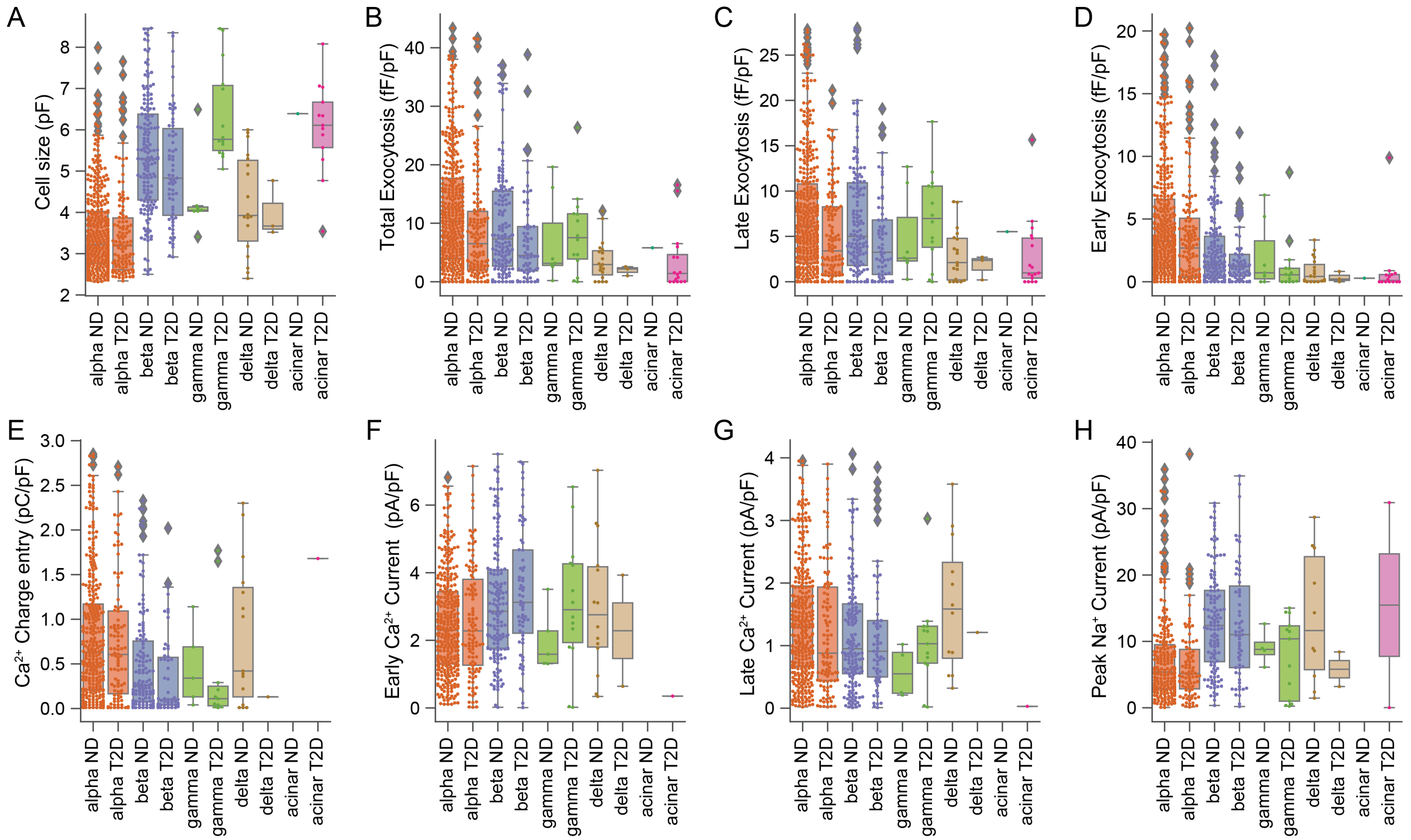


**Supp. Figure 2**

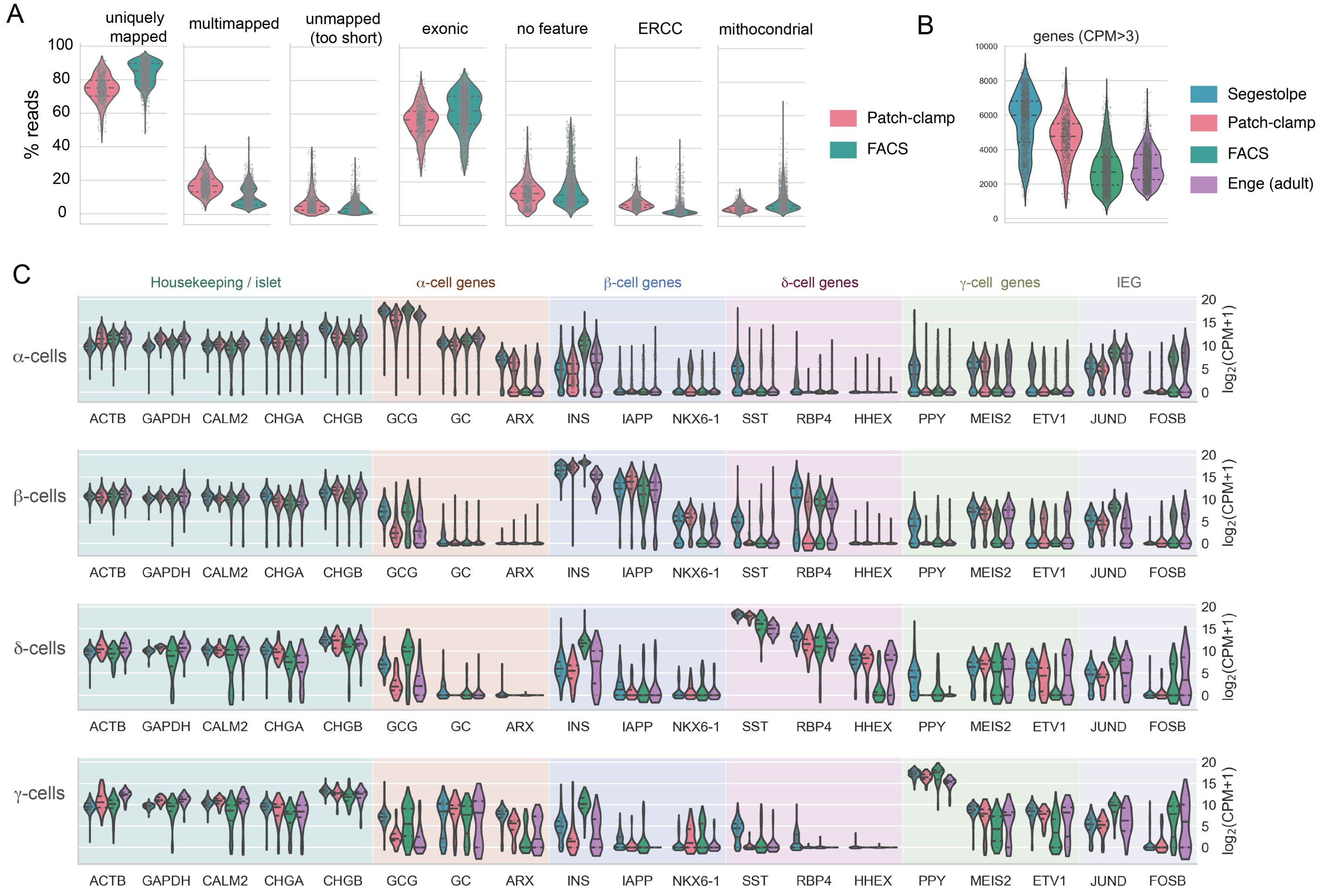


**Supp. Figure 3**

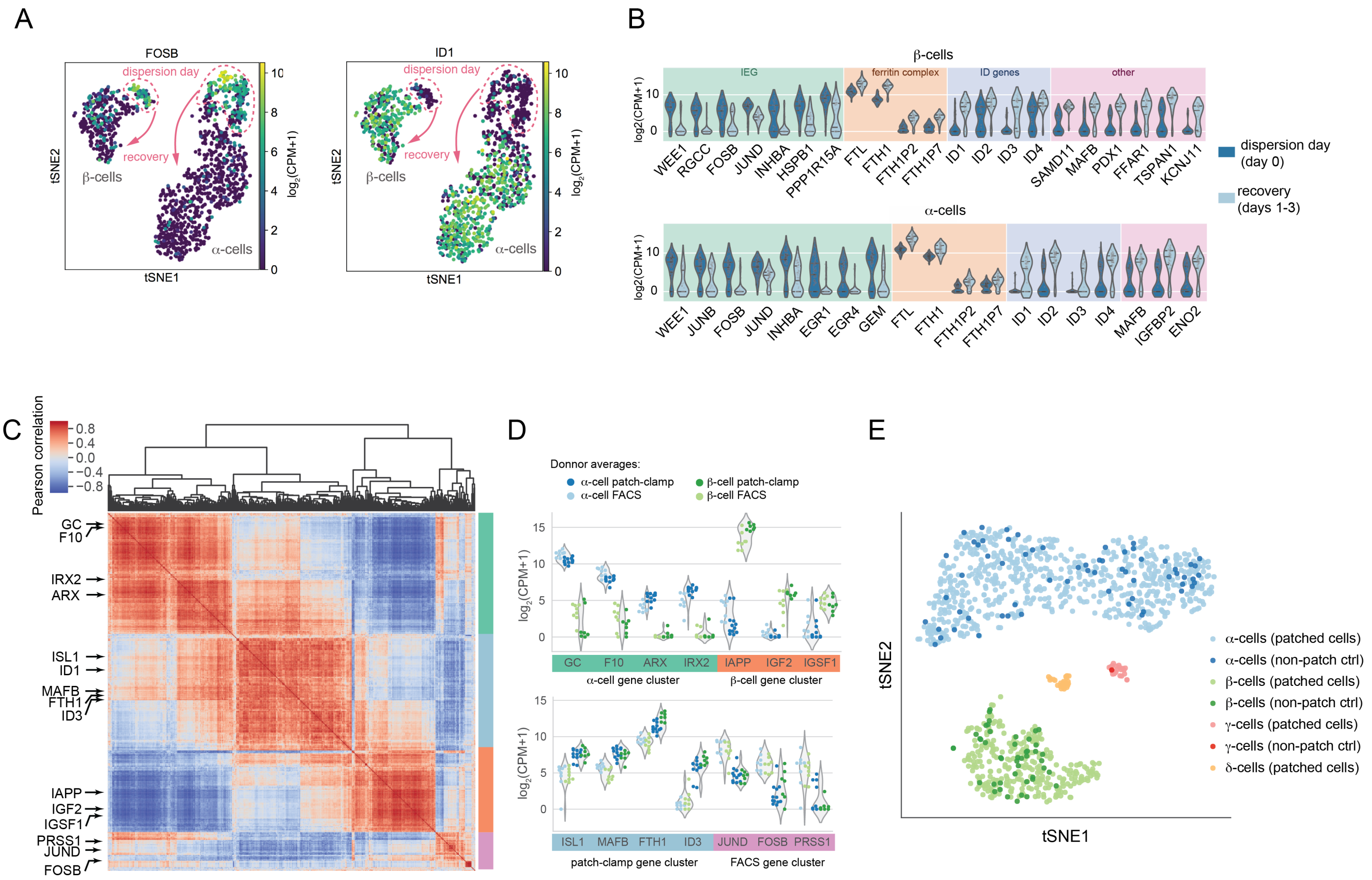


**Supp. Figure 4**

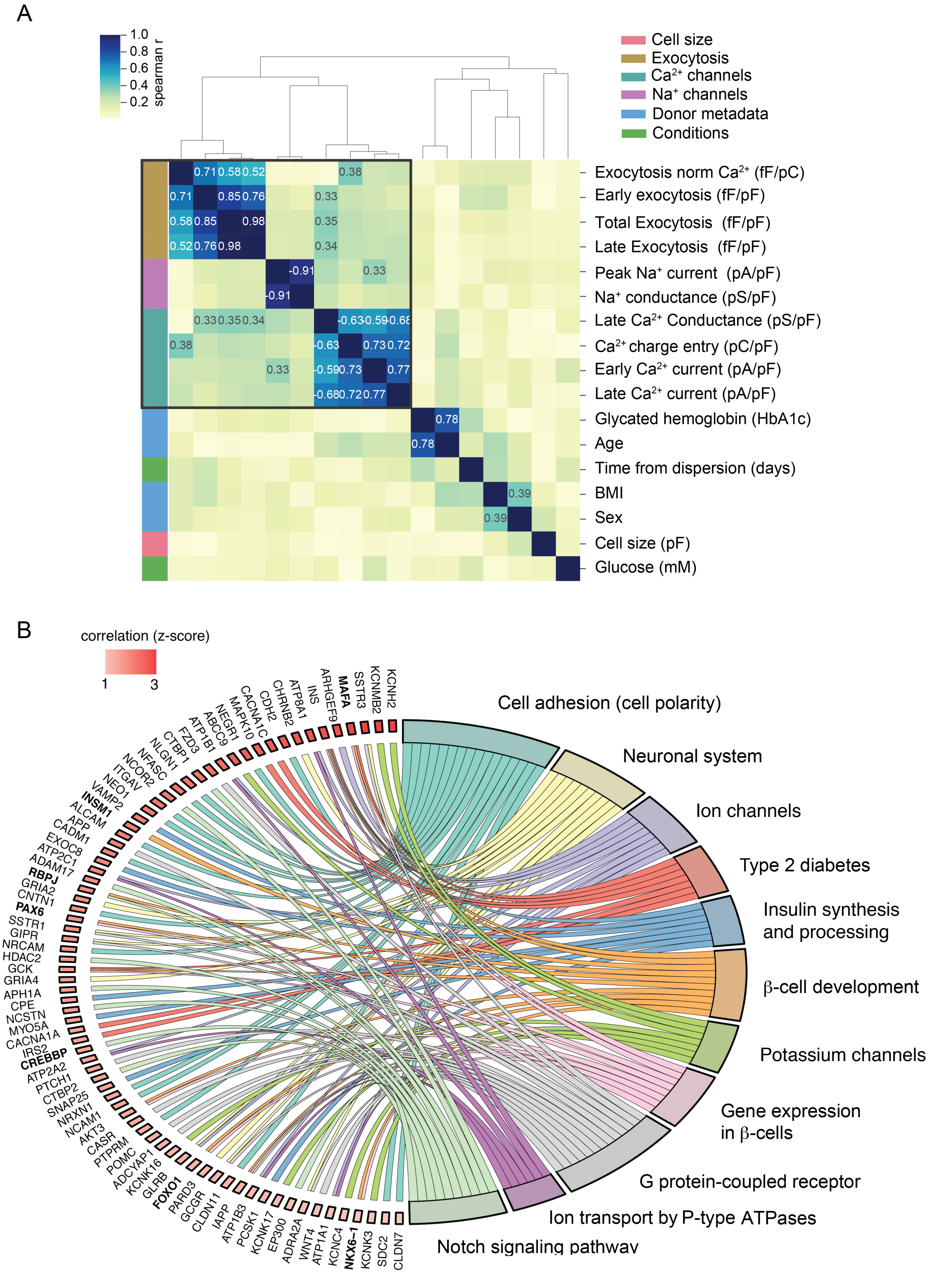


**Supp. Figure 5**

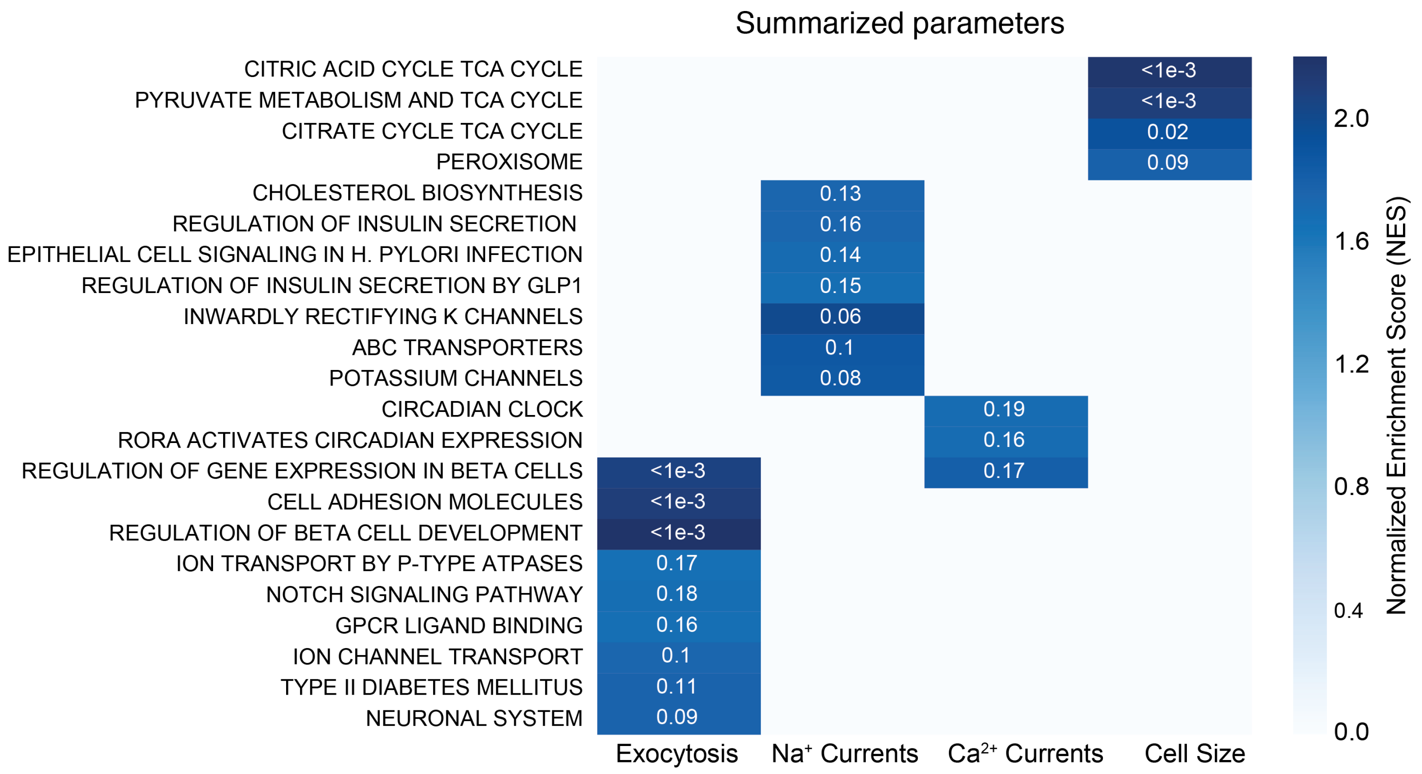


**Supp. Figure 6**

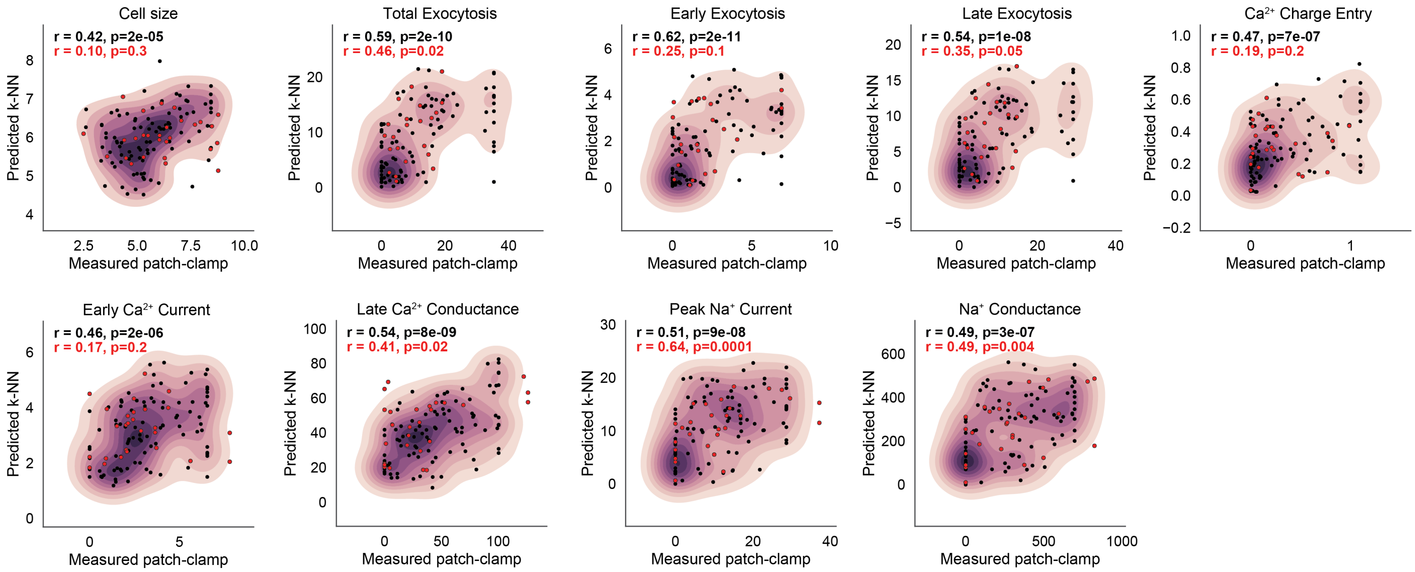


**Supp. Figure 7**

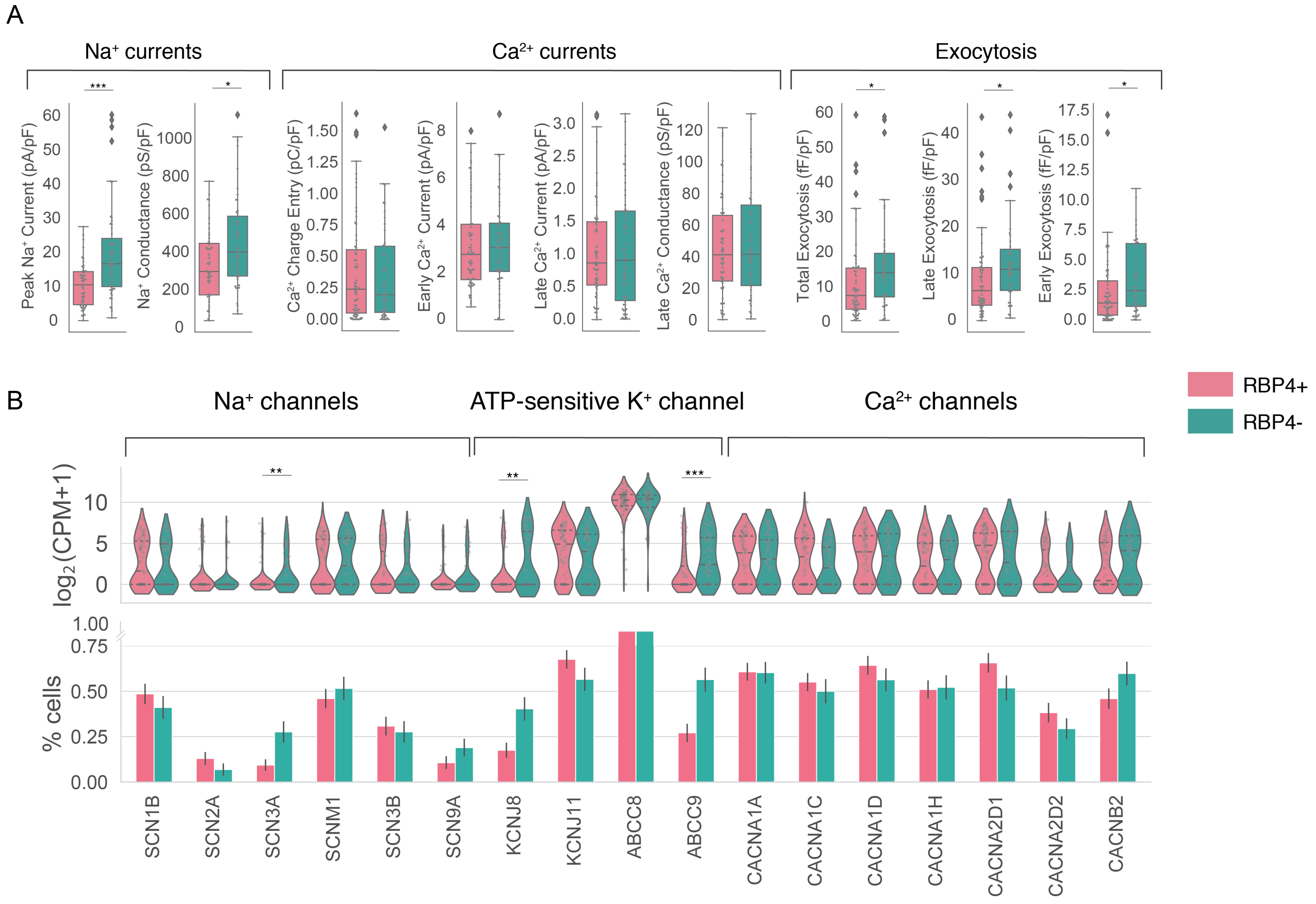


**Supp. Figure 8**

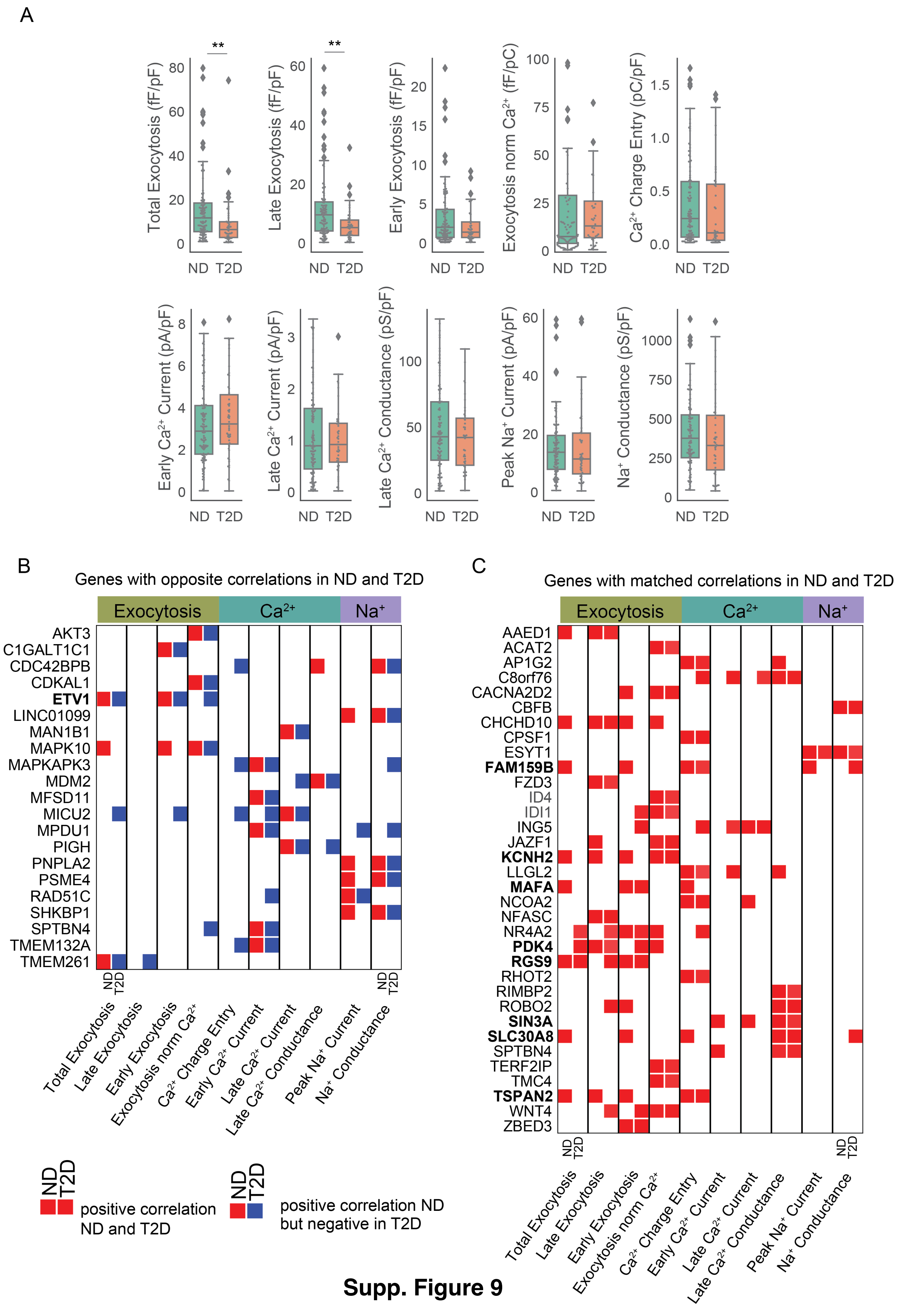


**Supp. Figure 9**

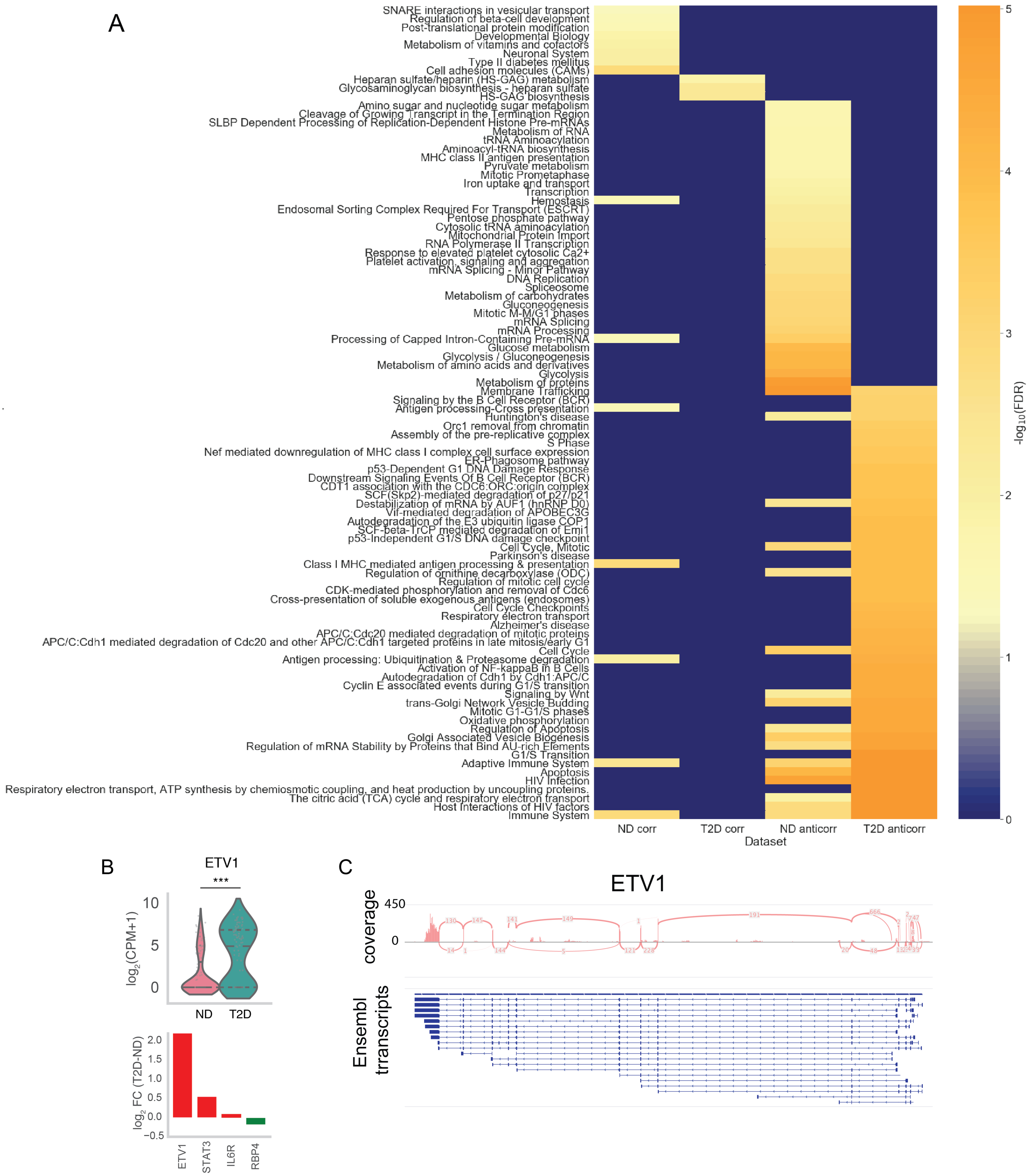


**Supp. Figure 10**

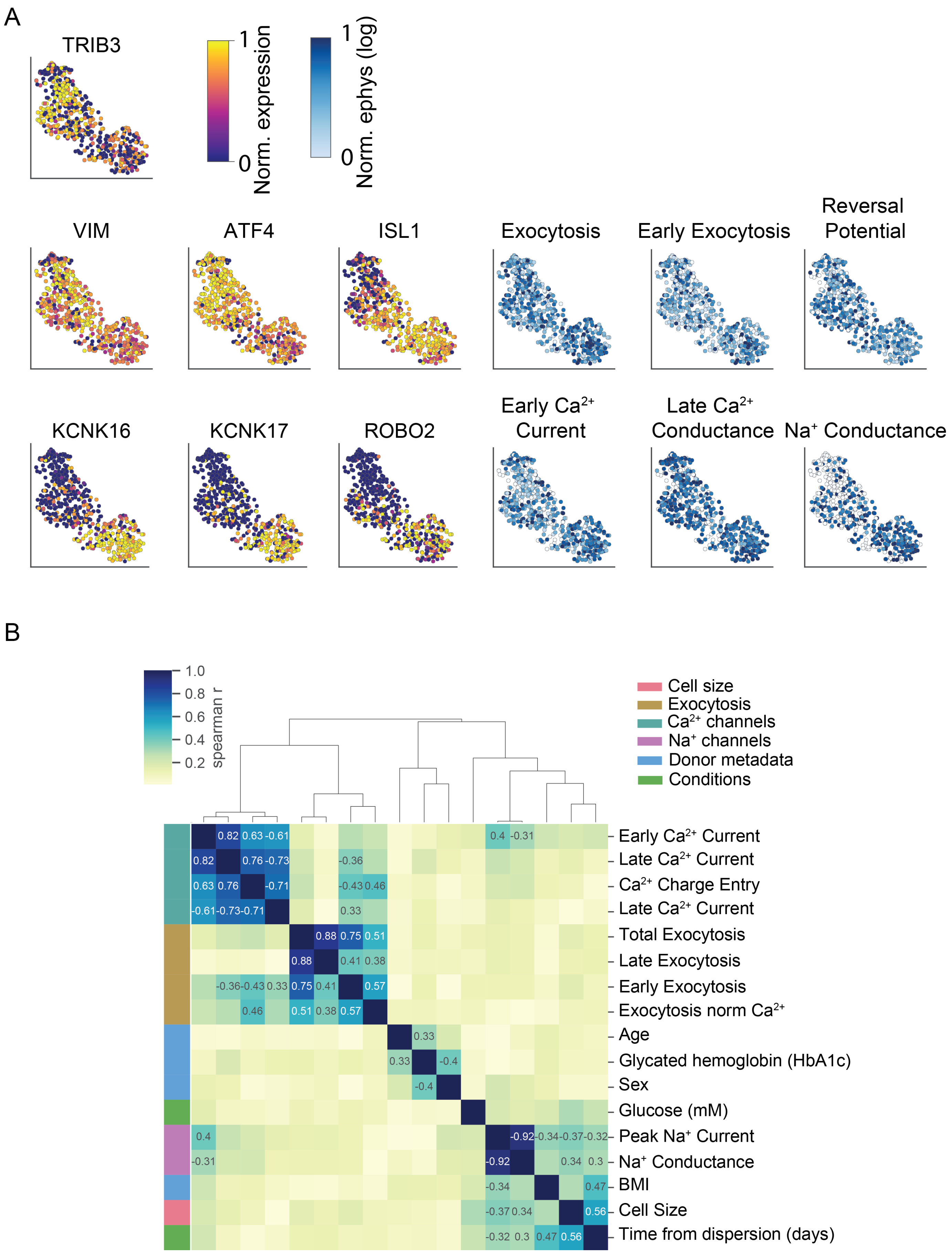


**Supp. Figure 11**

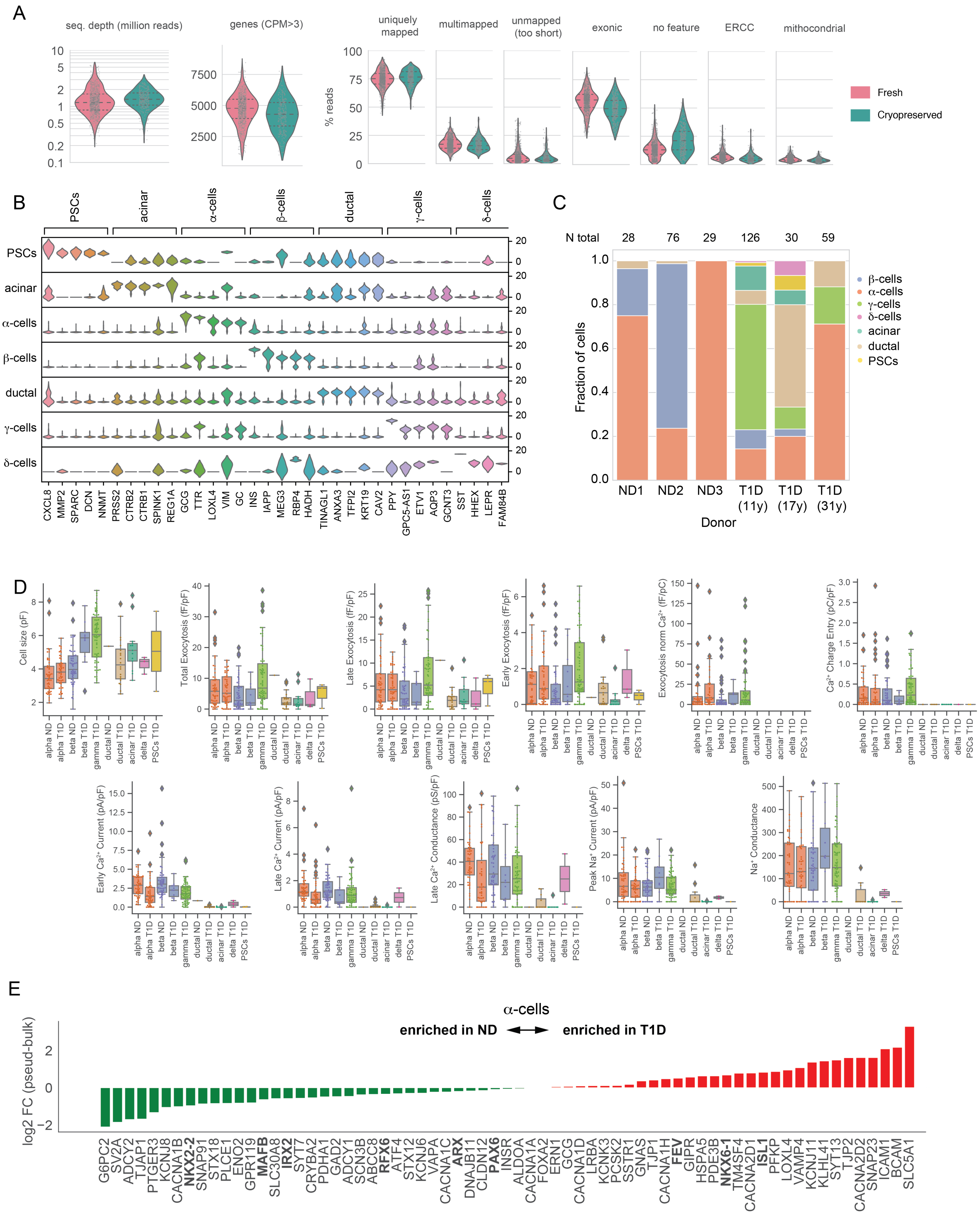


**Supp. Figure 12**
